# The cryoEM structure of the *Hendra henipavirus* nucleoprotein reveals insights into paramyxoviral nucleocapsid architectures

**DOI:** 10.1101/2023.12.07.570572

**Authors:** Tim C Passchier, Joshua B R White, Daniel P Maskell, Matthew J Byrne, Neil A Ranson, Thomas A Edwards, John N Barr

**Author notes:** Current address: Department of Biology, University of York, York, YO10 5DD, UK. Current address: Exscientia, The Schrödinger Building Oxford Science Park, Oxford UK, OX4 4GE. Current address: Larkin University, College of Biomedical Sciences, 18301 N Miami Avenue, Miami, FL 33169 USA. Corresponding authors: Thomas A Edwards; John N Barr.

## Abstract

We report the first cryoEM structure of the *Hendra henipavirus* nucleoprotein in complex with RNA, at 3.5 Å resolution, derived from single particle analysis of homotetradecameric RNA-bound N protein rings exhibiting D14 symmetry. The structure of the HeV N protein adopts the common bi-lobed paramyxoviral N protein fold; the N-terminal and C-terminal globular domains are bisected by an RNA binding cleft containing six RNA nucleotides and are flanked by the N-terminal and C-terminal arms, respectively. In common with other paramyxoviral nucleocapsids, the lateral interface between adjacent N and N_+1_ protomers involves electrostatic and hydrophobic interactions mediated primarily through the N-terminal arm and globular domains with minor contribution from the C-terminal arm. However, the HeV N multimeric assembly uniquely identifies an additional interaction between N_+1_ and N_-1_ protomers. The model presented here broadens the understanding of RNA-bound paramyxoviral nucleocapsid architectures and provides a platform for further insight into the molecular biology of HeV, as well as the development of antiviral interventions.

## Introduction

The non-segmented, negative-sense RNA virus *Hendra henipavirus* (HeV) emerged in 1994 as a fatal infectious agent on the Hendra racetracks of Brisbane, Australia, leading to the deaths of 15 horses and one human [1,2]. Since its discovery, there have been over 65 recognised and suspected outbreaks of HeV along the Australian east coast, with documented annual outbreaks since 2006 [3]. Serological sampling of wild populations has identified pteropid fruit bats as the natural reservoir of HeV [4,5]. While all recorded spillover events of HeV have occurred in Australia, the natural range of pteropid bats extends much further north into Asia, potentially extending the at-risk area [6]. Indeed, sampling among bats and humans in Africa has uncovered neutralising antibodies in both populations against HeV, as well as fellow genus members *Nipah henipavirus* (NiV) and *Ghanaian bat henipavirus* (GhV) [7]. Seropositivity in humans was most common amongst individuals dealing in bat bush meat or inhabiting areas of active deforestation [7], highlighting the increasing spillover risk of emerging zoonotic viruses due to human encroachment on natural landscapes. HeV infection typically exhibits a predominantly respiratory and neurological tropism, explaining the common symptoms of HeV infection including pulmonary haemorrhage and oedema, encephalitis, and meningitis and carries a mortality rate of 60-75% for both equine and human victims [8,9]. Disease progression is generally rapid and most infected individuals die within days, or in the case of horses are euthanised. However, one human case of recovery and subsequent fatal relapse 14 months after infection has been reported that resembles the brain inflammation seen in persistent *Measles morbillivirus* (MeV) infection [10]. Documented HeV transmission is predominantly through contact with bodily fluids and the data reported on spillover events suggests that inter-equine transmission can occur [1,2]. No direct transmission from bats to humans, or humans to humans, has yet been described, with all human cases the result of contact with infected horses. However, highly productive inter-human transmission of the closely related NiV has been observed in an outbreak with a mortality rate of around 75%, suggesting that similar outbreaks of HeV with inter-human transmission may occur in the future [11].

The HeV negative-sense single-stranded RNA genome is 18,234 nucleotides long, encodes six genes, and conforms to the paramyxoviral ‘rule of six’ [12–14]. The first gene encodes the nucleoprotein (N protein) which associates with the viral genomic and antigenomic RNAs to form the helical nucleocapsid [14–16]. The nucleocapsid associates with the viral RNA dependent RNA polymerase (RdRp) comprising the large (L) and phospho (P) proteins to form the holonucleocapsid. The N-RNA interaction is critical for virus multiplication as genomic RNA is only recognised by the RdRp as a template for mRNA transcription and genome replication when in the context of the assembled nucleocapsid (reviewed in [17]), and incorporation of genomic RNA into new virions depends on interactions between N protein within the nucleocapsid and the viral matrix (M) protein [17–19].

Paramyxoviral nucleocapsids, as well as those from the closely related family *Pneumoviridae*, classically exhibit a flexible, hollow, left-handed helical architecture under electron microscopy (EM), showing a characteristic herringbone pattern on transmission micrographs [20–26]. In these nucleocapsids, neighbouring N protomers laterally interact through their globular domains and completely encapsidate the viral genomic and antigenomic RNA within a continuous RNA binding cleft [27–29]. The N protomer of interest is topographically referred to as the index nucleoprotein (N_i_) while the immediately adjacent protomers upstream and downstream of the N_i_ are referred to as index +1 N protein (N_i+1_) or index −1 N protein (N_i-1_), respectively. Each paramyxovirus N protomer covers exactly six RNA bases, which obeys the paramyxovirus ‘rule of six’, with the RNA bases generally following a ‘three in, three out’ binding register where three RNA bases are oriented in towards the RNA binding cleft and three RNA bases are oriented out towards the solvent [12,13,29–31]. Each helical turn of the paramyxoviral nucleocapsid comprises approximately 13 N protomers, although the exact number differs between viral species and is not necessarily an integer [21,26–29,31,32].

In addition to the extended helical nucleocapsid architecture, other atypical nucleocapsid assemblies have been observed in recent years. The formation of N protein-RNA ring complexes has been described for recombinantly-expressed nucleoproteins from various viruses, such as decameric N-RNA rings of *Human orthopneumovirus* (RSV) and *Vesiculovirus indiana* [25,33], undecameric N-RNA rings of *Lyssavirus rabies* [34], and tridecameric N-RNA rings of *Mammalian orthorubulavirus 5* [35]. Interestingly, features that resemble N-RNA rings were also observed packaged within RSV virions [36]. Furthermore, so-called ‘clam-shell’ nucleocapsid assemblies have been observed from recombinantly expressed nucleoproteins [32,37,38]. These clam-shells are formed by two helical nucleocapsids engaged in back-to-back stacking and have been found for *Avian orthoavulavirus 1* (formerly Newcastle disease virus; NDV), *Murine respirovirus*, and NiV [32,37,38].

Structural information on the HeV N protein and nucleocapsid assembly is scarce, collectively no more than a handful of negative-stain EM micrographs of isolated HeV nucleocapsids [39] or recombinantly expressed HeV nucleocapsid-like filaments [40] exist. While these confirm the similar herringbone morphology of HeV nucleocapsids to those of closely related viruses, a high-resolution reconstruction of the HeV N protein and nucleocapsid assembly has not yet been determined. Here, we report for the first time the HeV N protein structure in the context of a double-ring assembly as determined through cryoEM from recombinantly expressed HeV N protein. The approach taken reveals the full HeV Ncore structure (residues 1-394) and further elucidates established and novel paramyxoviral N-N and N-RNA interactions.

## Results

### Architecture of the HeV N protein-RNA assembly

HeV N protein was recombinantly expressed in *E. coli* and purified through IMAC using an N-terminally fused hexahistidine-SUMO affinity purification tag, followed by proteolytic removal of the affinity purification tag and subsequent SEC purification (Supplementary Figure S1). The purified HeV N sample was applied to cryo-grids and automated data collection was performed, including on-the-fly (OTF) pre-processing of the 3548 collected micrographs (Supplementary Figure S2) [41]. The micrographs contained a range of particles exhibiting circular and squat ring morphologies. Selection of these particles followed by 2D and 3D classification in RELION [42] revealed that these morphologies represent different orientations of the same multimeric N protein assembly, viewed across a 14-fold and 2-fold axis of symmetry, respectively (Supplementary Figure S3). Subsequent 3D refinement of a curated particle stack comprising 5638 particles and using D14 symmetry resulted in a 3.5 Å resolution EM density map, revealing a previously uncharacterised multimeric nucleoprotein assembly (Supplementary Figure S3, Supplementary Figure S4). The final map is a dimer of tetradecameric rings which enabled model building resulting in the first reported HeV N protein structure (Figure 1, Supplementary Figure S3, Supplementary Figure S4, Supplementary Table S1). In the HeV N protein assembly, two rings stack ‘bottom-to-bottom’, where the globular N-terminal domains (NTDs) intercalate to form a dimer of tetradecamers (Figure 1a-c). Each tetradecameric ring is topped with the C-terminal arms (Ct-arms) which cover parts of the globular C terminal domains (CTDs) (Figure 1a-c). The N-terminal arm (Nt-arm) lies completely within the particle’s central cavity where it crosses over the NTD and interfaces with the back face of the CTD (Figure 1b).

**Figure 1.**
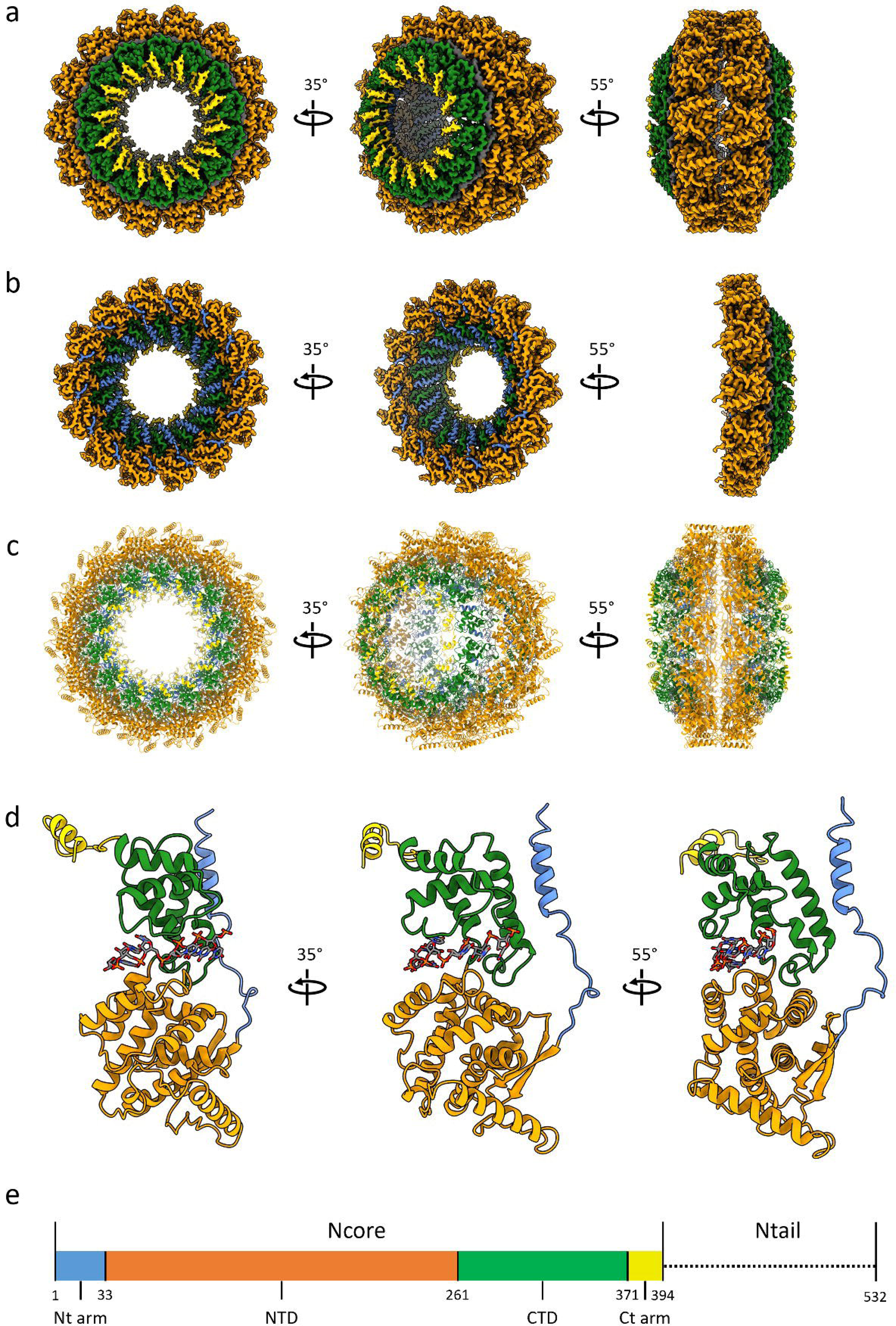
Structure of the HeV N protein-RNA double-ring assembly. The cryoEM map (**a-b**), resolved to 3.5 Å resolution, and reconstructed HeV N multimer (**c**) and protomer (**d**) were coloured according to HeV N protein domains (**e**). Views of the map showing a complete double-ring assembly (**a**) or single tetradecameric ring (**b**) are shown aligned to the corresponding view of the reconstructed multimer (**c**). The multimeric assembly reveals that the top of each tetradecameric ring consists of the C-terminal arm (Ct-arm; gold) which lies on top of the globular C-terminal domain (CTD; green) (**a**). The globular N-terminal domains (NTDs; orange) intercalate to form a dimer of tetradecamers (right panels). The N-terminal arm (Nt-arm; blue) lies completely within the double-ring central cavity (**b**) where its loop region jumps over the NTD and its alpha-helical coil interfaces with the CTD. **d**) A focused view of a single HeV N protomer from the double-ring assembly shows the bound RNA (sticks; coloured by element). **e**) Schematic, to-scale linear representation of the HeV N protein domain architecture indicating the Ncore and Ntail regions. The Ncore consists of the NTD and CTD, each of which are flanked by a flexible arm; the Nt-arm and the Ct-arm. The intrinsically disordered Ntail region was not resolved in the cryoEM map nor modelled in the reconstruction.

### Structure of the HeV N protein

The EM map of the HeV N protein assembly fully resolved the N protein core (Ncore), however no density was observed in the map corresponding to the intrinsically disordered C-terminal N protein tail (Ntail, Ala395-Val532). The HeV Ncore structure follows the common bi-lobed paramyxoviral N protein fold and includes the Nt-arm (Met1-Thr32), NTD (Thr33-Glu260), CTD (Glu261-His370), and Ct-arm (His371-Ala394), as well as bound *E. coli* host-derived RNA (Figure 1d-e). Both the Nt- and Ct-arm are linked to the globular domains by flexible loops and each forms an alpha helix in proximity to the CTD (Figure 1d). The N protein surface area is overall hydrophilic, with three prominent hydrophobic patches that are covered by neighbouring N protomers in the assembly (Supplementary Figure S5, Supplementary Figure S6). The Nt-arm alpha helix (α1) contains a strongly negatively charged, solvent-exposed region facing the central cavity consisting of residues Asp3, Asp6, and Glu7 (Figure 2).

**Figure 2.**
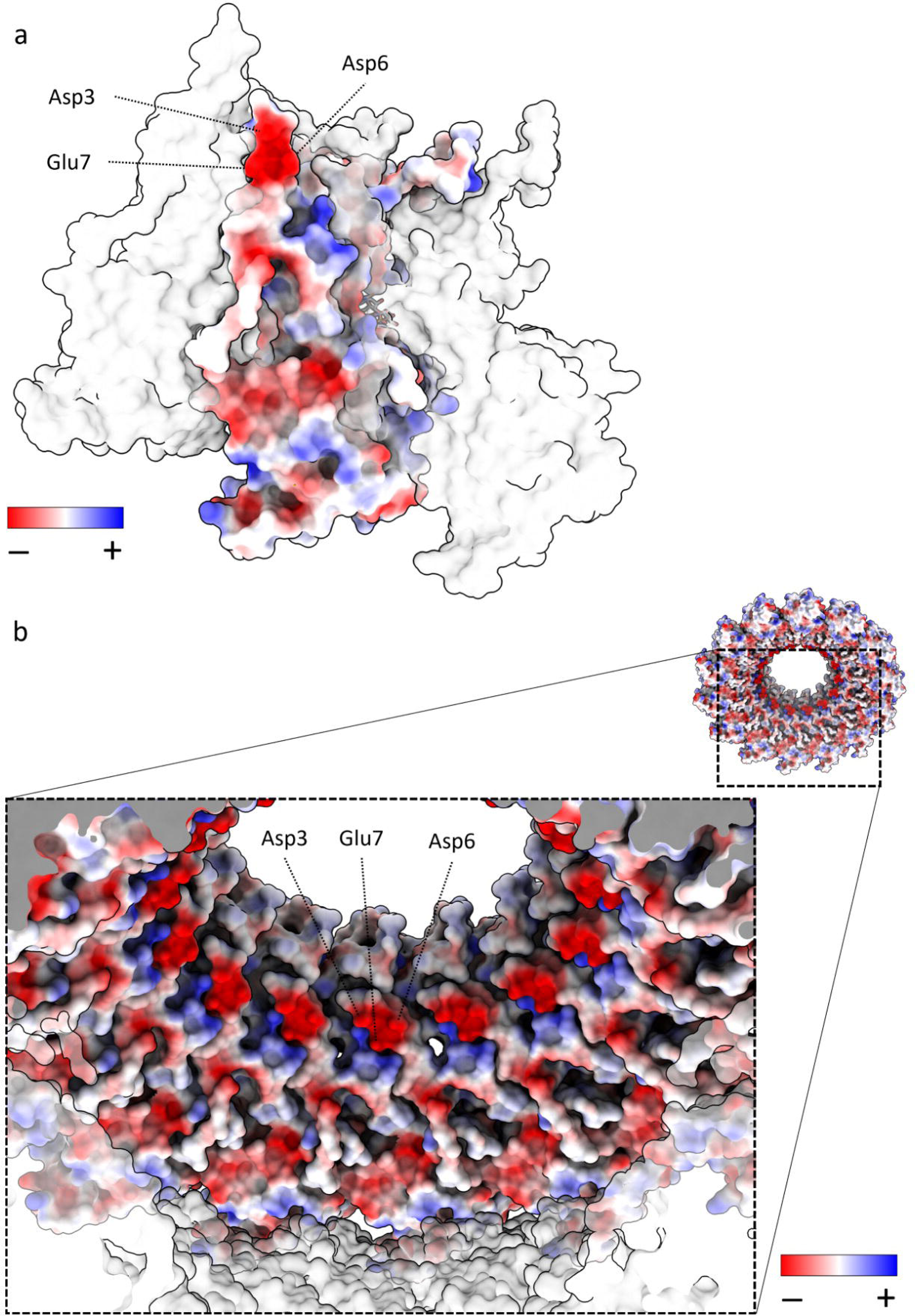
HeV N protein double-ring surface values. The presence of a strongly negatively charged patch (Asp3, Asp6, Glu7) is indicated within the context of a HeV N protomer (**a**) or a tetradecameric ring (**b**). The protomer or ring of interest is coloured according to electrostatic surface potentials for negative (red, -) and positive (blue, +) residues or uncoloured (white) for neutral residues. HeV N protomers or tetradecameric ring adjacent to the protomer or ring of interest, respectively, are transparent.

### HeV N – N protein interactions in the multimeric assembly

The HeV N protein double-ring assembly is formed through extensive lateral interactions in each tetradecameric ring as well as intercalating interactions between two tetradecameric rings. The lateral interfaces consist of electrostatic and hydrophobic interactions and are mediated primarily through the Nt-arms, NTDs, and CTDs of two neighbouring N protomers, with contributions from the Ct-arm, over a total buried surface area of ∼2930 Å^2^, calculated via PDBe PISA [43] (Supplementary Figure S5, Supplementary Figure S6). At a calculated buried surface area of ∼1195 Å^2^, the N_i_ - N_i+1_ interface mediated by globular domains represents just over a third of the total buried surface area between the neighbouring protomers. The globular domain interface consists of extensive electrostatic interactions, with the N_i_ surface containing mostly positively charged patches and the N_i+1_ surface mostly negatively charged patches (Supplementary Figure S5). While the contribution of the Ct-arm is minor, with a calculated buried surface area of only ∼340 Å^2^, the Nt-arm contributes extensively to the lateral interactions. At a calculated buried surface area of ∼1740 Å^2^, this interface represents just over half of the total lateral interface between N_i_ and N_i+1_. Aside from electrostatic patches, a conserved hydrophobic region consisting of a triad of aromatic residues (Phe267, Phe268, and Tyr301) in the CTD of the N_i_ protomer and Phe11 in α1 of Nt-arm N_i+1_ protomer forms a key interface [28,32] (Figure 3a). A further hydrophobic cleft in the N_i_ NTD (Tyr80, Ser81, Leu87, Val162, Val165, Ile166, Ile234, and Val249) is occupied by the N_i+1_ Nt-arm loop Ala27-Thr33. Within this N_i+1_ Nt-arm loop, residue Leu31 is buried inside a pocket in the N_i_ NTD consisting of Asn43, Arg48, Tyr80, Glu82, Met158, Leu161, Val162, and Val165 (Figure 3b). Beyond the extensive N_i+1_ Nt-arm interface with the Ni NTD and CTD, a minor contact exists between the utmost amino-terminal residues of the N_i+1_ Nt-arm (Met1, Ser2, Ile4, and Phe5) and the N_i-1_ Ct-arm loop region Gln376-Lys381 (Figure 4). This N_+1_ - N_-1_ interface represents a novel interaction that, to the best of our knowledge, has not been described in the literature for paramyxoviral nucleocapsids and will be referred to as the ‘elbow interface’. Since the elbow interface only has a calculated buried surface area of ∼80 Å^2^, it is unlikely to contribute strongly to N-N interactions. A defining feature of this HeV N protein double-ring assembly is the gear-wheel interface between the two tetradecameric rings. This intercalation zone consists solely of residues in the NTDs of N protomers in the top ring and N protomers in the bottom ring (indicated by the lower-case n) (Figure 5). The N_i_ protomer contacts two protomers in the bottom ring (n_i_ and n_i+1_), with a total calculated buried surface area of ∼545 Å^2^. A clear, reciprocal electrostatic interface exists between N_i_ and n_i+1_ with the residues Lys120, Glu123, and Glu124 where the two Glu residues in N_i_ form a tab that interacts with the Lys residue in n_i+1_ and *vice versa* (Figure 5b). The poor electron density surrounding loop Met114-Asp119 forced the stubbing of side chains and, accordingly, these were modelled as alanines (indicated in italics) (Figure 5b-c). An electrostatic interaction may exist between the N_i_ Asp157 and n_i_ *Arg116*, as they are in proximity and there is ample space for the Arg side chain. Similarly, N_i_ residues Ser44 and Glu46 form a patch that may interact with n_i_ residues *Met114* and *Glu115* which are in close proximity. The absence of clear electron density neither supports nor refutes these putative interactions and improved local resolution may further elucidate the intercalation zone and increase the calculated buried surface area.

**Figure 3.**
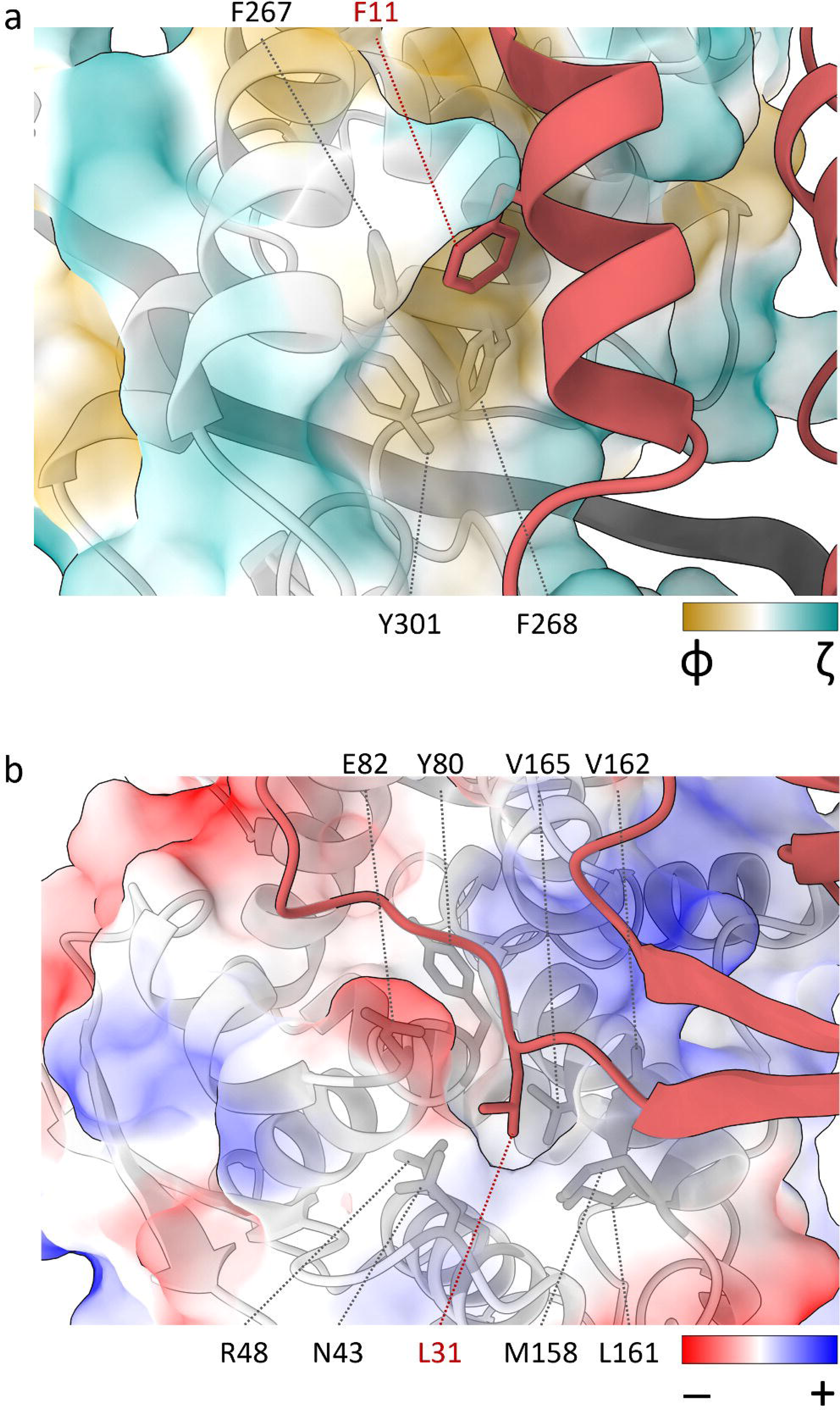
HeV N_i+1_ Nt arm residues Phe11 and Leu31 are buried in conserved pockets on the neighbouring N_i_ globular domains. **a**) The N_i+1_ Nt-arm residue Phe11 (red) sits within a conserved hydrophobic pocket consisting of the neighbouring N_i_ protomer CTD residues Phe267, Phe268, and Tyr301 (grey sticks). Surface colours are mapped to Kyte-Doolittle hydrophobicity scale [88] from most hydrophilic (green, ζ) to most hydrophobic (gold, φ). The RNA is represented as a dark grey ribbon. **b**) The N_i+1_ Nt-arm residue Leu31 (red) engages in electrostatic interactions with the neighbouring N_i_ protomer NTD residues Asn43, Arg48, Tyr80, Glu82, Met158, Leu161, Val162, and Val165 (grey sticks). Electrostatic surface potentials are coloured for negative (red, -) and positive (blue, +) residues or uncoloured (white) for neutral residues.

**Figure 4.**
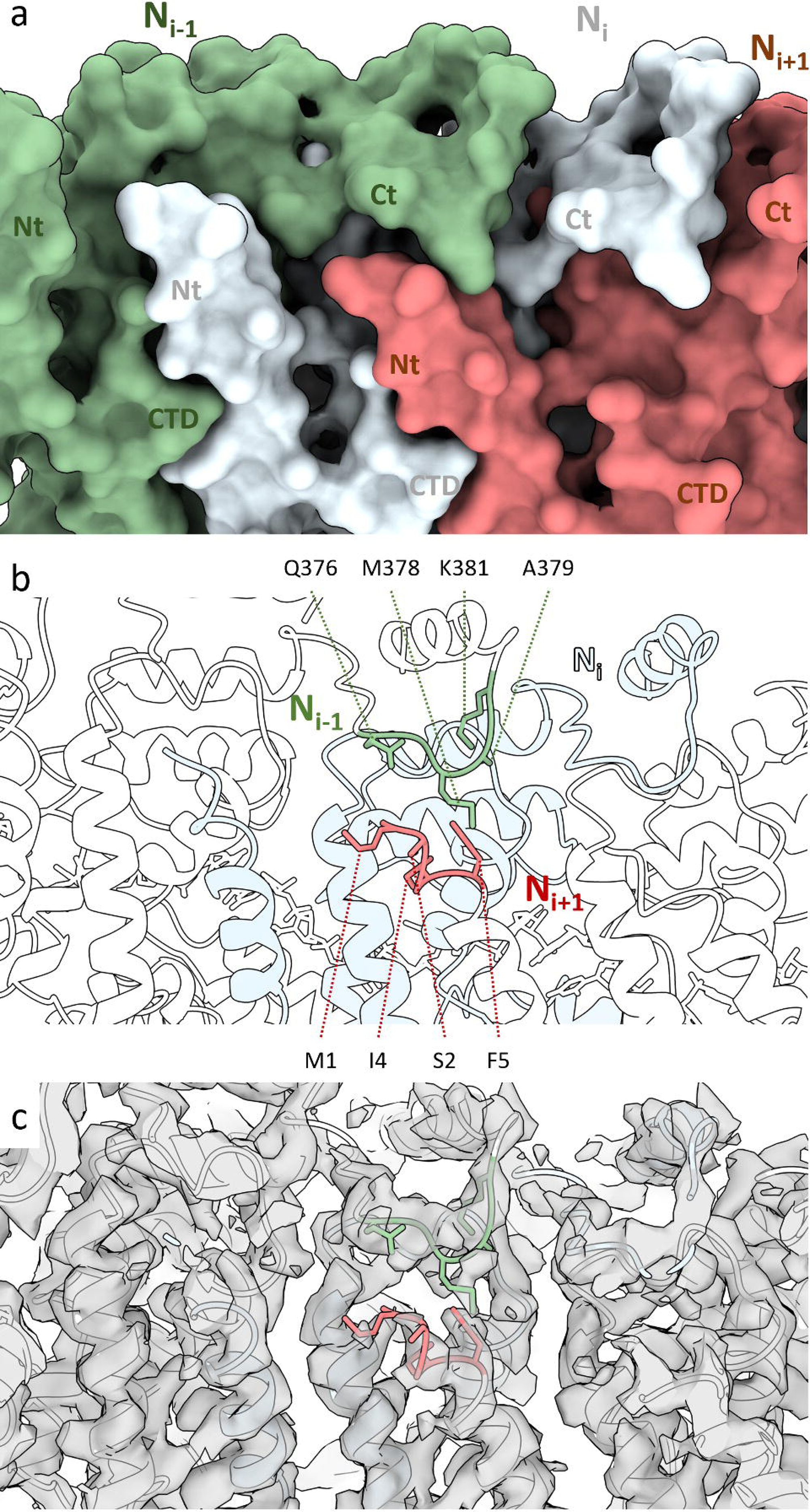
The HeV double-ring assembly forms elbow interactions bridging non-neighbouring protomers. Three N protomers in the assembled ring, showing the N_i_ (pale blue), N_i+1_ (coral), and N_i-1_ (green) protomers, centred on the elbow interaction between the N_i+1_ Nt-arm and the N_i-1_ Ct-arm. **a**) A surface render of three adjacent N protomers within the assembled ring with their Nt-arm (Nt), CTD, and Ct-arm (Ct) indicated. **b-c**) A ribbon visualisation of three N protomers. Protomers N_i+1_ and N_i-1_ are coloured white and the N_i_ is coloured pale blue. Putative interacting residues are coloured coral and green and presented as sticks. **b**) Residues putatively involved in the elbow interactions are shown as sticks and indicated. **c**) The cryoEM double-ring density map is shown as a transparent overlay.

**Figure 5.**
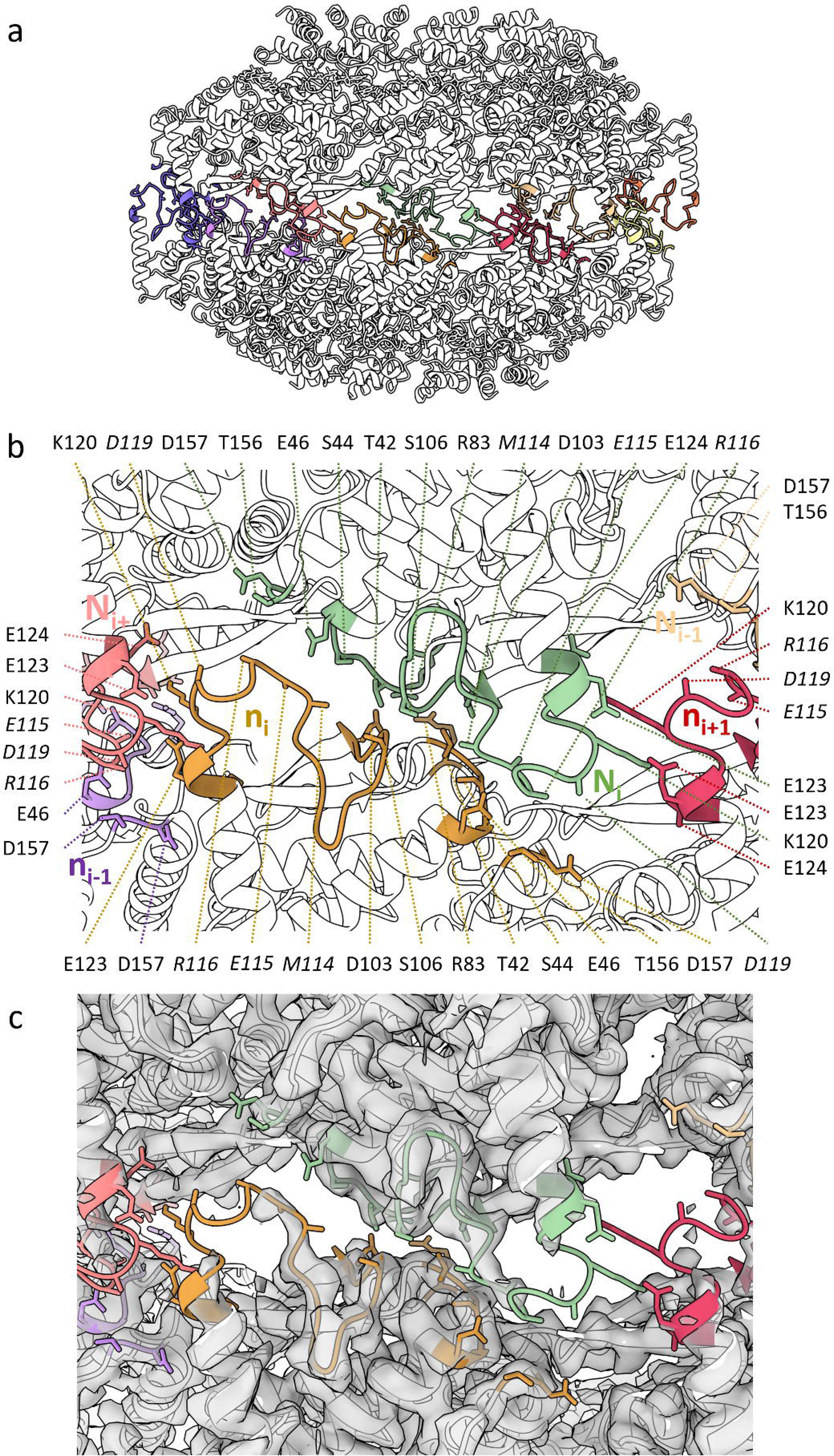
The HeV double-ring assembly intercalation zone is poorly resolved. A ribbon visualisation of a dimer of N pentamers. N protomers are coloured white with loops belonging to the intercalation zone coloured per protomer. Putative interacting residues are presented as sticks. **a**) An overview of the dimer of pentamers assembly showing the intercalation zone. **b-c**) The intercalation zone centred around the N_i_ protomer (lime) of the top ring, showing the N_i+1_ (pink) protomer as well as the bottom ring protomers N_i_ (n_i_; gold), N_i-1_ (n_i-1_; purple), and N_i+1_ (n_i-1_; coral). **b**) Residues putatively involved in the intercalation interactions are shown as sticks and indicated. Italicised residues are stubbed and modelled as alanines. **c**) The cryoEM double-ring density map is shown as a transparent overlay revealing areas of poor density.

### HeV N – RNA interactions

The HeV N assembly map showed clear density for single-stranded RNA in the RNA-binding cleft that separates the NTD and CTD, which was modelled as a poly-uridine; one hexa-uridine per N protomer (Figure 6, Supplementary Figure S7). The RNA binding cleft contains hydrophobic patches and is mostly positively charged near the RNA backbone and negatively charged near the RNA bases (Supplementary Figure S5). The RNA molecule is oriented in the conventional ‘three-base in, three-base out’ similar to other paramyxoviruses [28,29,32,35,37]. This orientation is facilitated by a 180° turn of the RNA backbone every three nucleotides. In the case of HeV N, the 3’-most three-base stack is oriented in towards the RNA-binding cleft while the 5’-most three-base stack are oriented towards the solvent (Figure 6, Supplementary Figure S7. The charged RNA-binding cleft consists of a cleft floor and two ridges that follow the orientation of the RNA chain (Figure 6a-c). Residue segments Thr181-Gln200, containing α8, in the NTD and the loop Ser344-Tyr354 in the CTD form the solvent-facing cleft ridges. The cleft floor consists of the NTD-CTD-connecting loop Glu260-Ala265 continuing into α12 until Arg272 as well as the α14-containing stretch Ser317-Gly326. Within the RNA-binding cleft, a series of conserved basic (Lys178, Arg192, Arg193, Arg352) and polar (Thr185, Gln199, Gln200, Tyr258, Gln319, Ser344, Tyr354) residues likely interact with the RNA chain through electrostatic and hydrophobic interactions (Figure 6b-d) [28,29,32,35,37].

**Figure 6.**
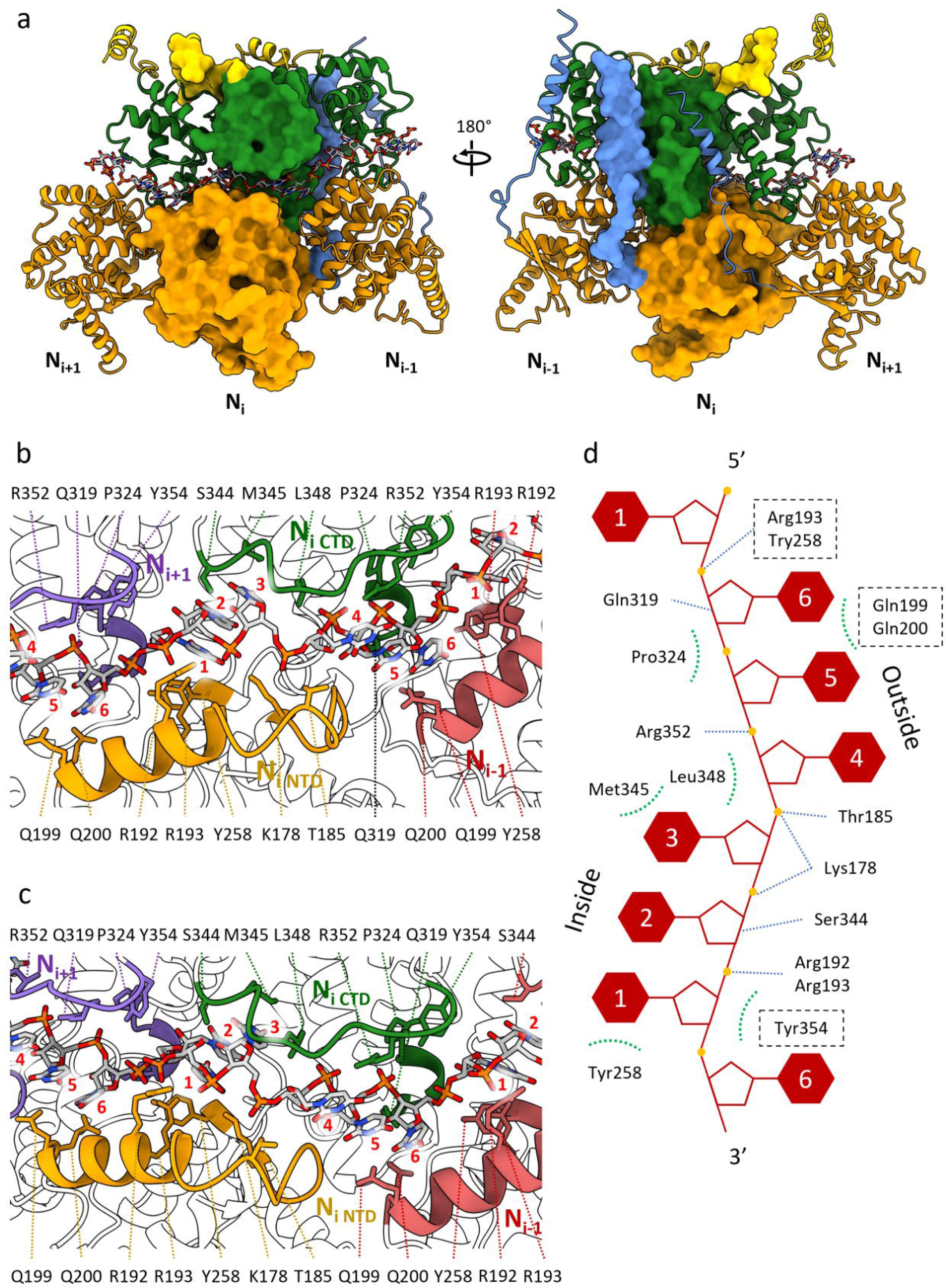
The HeV N protein binds RNA through electrostatic and hydrophobic interactions. **a**) Three N protomers within the assembled ring with the N_i_ protomer as a surface render and the neighbouring N_i+1_ and N_i-1_ protomers as ribbons. The N protein domains are individually coloured; the Nt-arm in blue, the NTD in orange, the CTD in green, and the Ct-arm in gold. The 18-mer RNA chain is visualised as sticks and follows heteroatom colouring. **b-c**) Two different close ups of the RNA binding cleft centred on the N_i_ protomer. N proteins are visualised as ribbons and the RNA chain as sticks. Helices and loops containing putative RNA interacting residues are coloured for the N_i_ (NTD; gold, CTD; green), the N_i+1_ (purple), and the N_i-1_ (coral). RNA bases in each hexamer are numbered 1 to 6 on a per-N protomer basis from the 3’-to the 5’-end. **d**) A topological schematic of the RNA-interacting residues centred around the Ni protomer and the corresponding RNA hexamer. RNA base stack orientations are shown and bases in each hexamer are numbered 1 to 6 from the 3’-to the 5’-end. Putative protein-RNA interactions are indicated for electrostatic interactions (blue dashed line) and hydrophobic interactions (green dashed arc). Residues belonging to neighbouring N protomers are bounded by a dashed box.

Adjacent to and contiguous with the RNA binding cleft floor is a large cavity surrounding the inward facing three-base stack flanked by the N_i_ NTD and CTD as well as α13 of the N_i+1_ CTD and α1 of the N_i+1_ Nt-arm (Figure 7). While the modelled RNA bases are all uridines, this cavity – here called the ‘in-base cavity’ – is sufficiently spacious to accommodate the larger purine bases and, correspondingly, N_i_ residues Tyr258, Glu260, Glu261, Phe268, Ser344, and Met345, and N_i+1_ residue Lys321 may be involved in interactions with these bases (Figure 7b).

**Figure 7.**
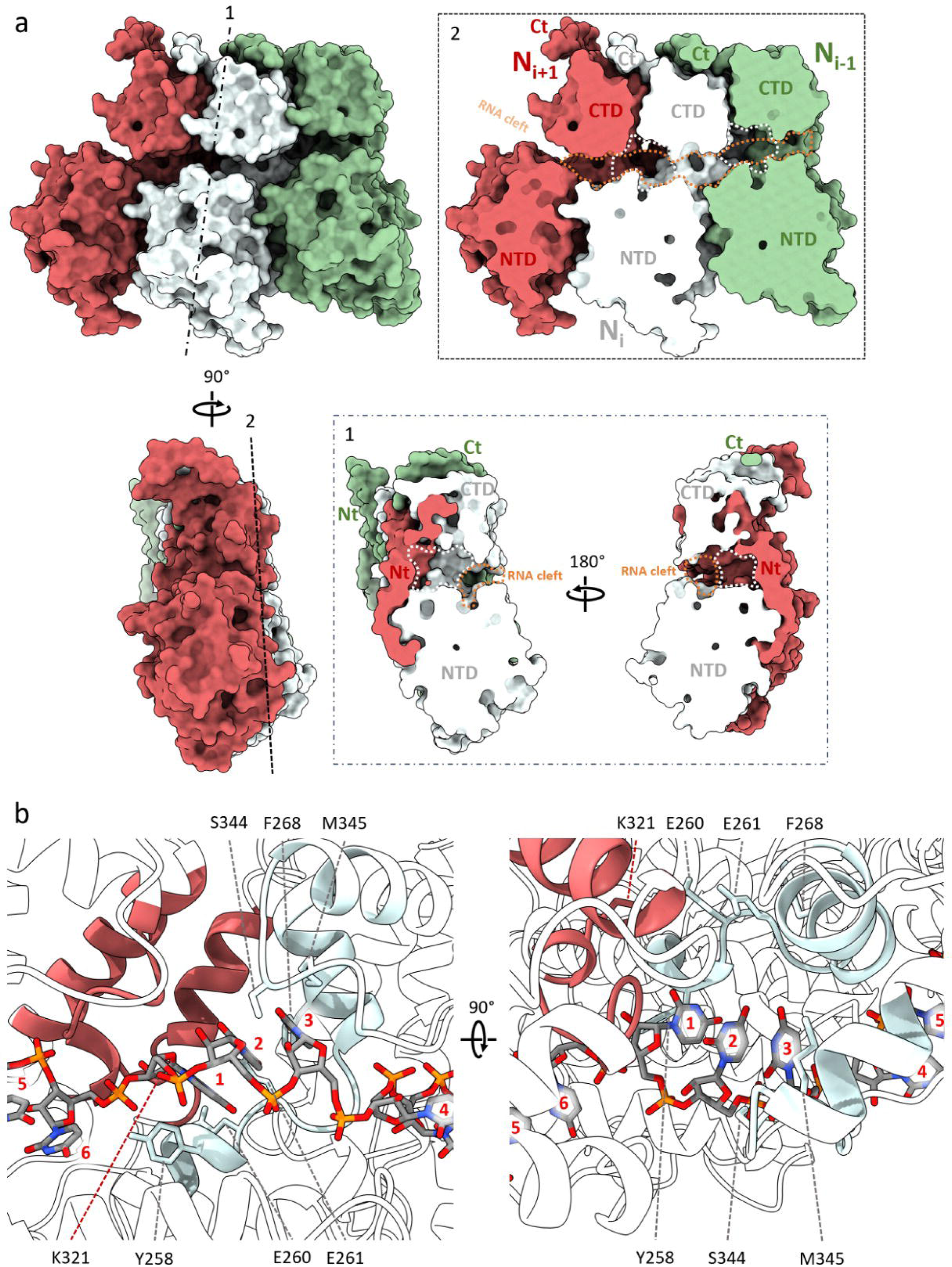
Neighbouring HeV N protomers form an in-base cavity. **a**) A trimer of N protomers as surface renders and individually coloured; N_i_ (light blue), N_i+1_ (coral), and N_i-1_ (green). Protomer domains (NTD, CTD, Nt-arm, and Ct-arm) are indicated. The trimer was sliced between two neighbouring protomers (1) and down the frontal face (2) to reveal the RNA binding cleft (orange dashed area) and in-base cavities (white dashed area). The RNA chain has been removed for clarity. **b**) An N protein dimer centred on the in-base cavity. Helices and loops that form the in-base cavity are coloured for the N_i_ (light blue) and the N_i+1_ (coral). Putative RNA-base interacting residue side chains are visualised as sticks and indicated. The N_i_ Ct-arm has been removed for clarity. RNA bases in each hexamer are numbered 1 to 6 on a per-N protomer basis from the 3’-to the 5’-end.

### Comparison with other paramyxoviral N protein assemblies

The HeV N protein fold resembles that of NiV N protein, its closest relative [8,14,32,44]. The NiV N protein structure has been solved recently through cryoEM [32] and earlier through X-ray crystallography [44]. Structural alignment of the HeV N protein model with the NiV N models reveals some key differences (Figure 8). The global RMSDs for the HeV N protein structure compared to that of the cryoEM-derived, RNA-bound NiV N protein structure and that of the NiV N protein structure solved through X-ray crystallography are very similar (0.756 Å and 0.783 Å, respectively). In the NiV X-ray structure, however, the NTDs and CTDs are rotated approximately 25° with respect to each other compared to the HeV N and NiV cryoEM structures [32] resulting in a shift of the CTD of up to 15.5 Å at its greatest extent (Figure 8a). This rotation in the crystal structure is likely induced by binding to the P protein Nt region which keeps the N monomers in an ‘open’ RNA-free conformation (N^0^) [16,44–47]. Three key differences appear from the alignment of HeV N with the two NiV N structures. Firstly, the HeV N Nt- and Ct-arms in the double-ring assembly are shifted compared to the Nt- and Ct-arms in the NiV N helical assembly (Figure 8b). Helical spiralling of a single split tetradecameric ring would generate a nucleocapsid-like helix and inevitably requires some minor conformational changes. The flexible nature of the Nt- and Ct-arm put them forward as suitable candidates to provide those changes. Secondly, the solvent facing NTD alpha-helix 5 and its succeeding loop (α5-loop) are also shifted compared to the homologous helix-loop in the NiV structure (Figure 8b). In fact, this helix-loop region exhibits significant structural and sequence variability among paramyxoviruses. Both its position and length are highly variable among the N proteins of closely related paramyxoviruses, including *Mammalian orthorubulavirus 5* (formerly human parainfluenza virus 5; PIV5), MeV, NDV, NiV, and SeV [29,32,35,37,38,44]. For example, the HeV N protein α5-loop region contains two amino acid substitutions compared with the NiV N protein; L108V and E137D, both of which are solvent exposed [32]. Whether there is a functional relevance to these substitutions is unknown, however in the context of a helical nucleocapsid the α5-loop region is the main solvent exposed section of the N protomer and may be important for interactions with the RdRp complex during transcription and replication, or could potentially interface in-phase with the M protein lattice during virion assembly, as proposed for other paramyxoviruses [17–19]. Furthermore, the α5-loop region has also been identified as the main interface in intercalating interactions in the HeV N protein assembly (Figure 5), as well as in clam-shell formation in NDV, NiV, and SeV nucleocapsid-like filaments (NLFs) [32,37,38]. The intercalating interface observed in the HeV N double-ring assembly suffered from local poorly resolved electron density (Supplementary Figure S4), which abrogated confident model building and forced the stubbing of residue side chains for *Met114*, *Glu115*, *Arg116*, *Arg117*, and *Asp119*. This poor local density may reflect the existence of weak interactions within the interface since stronger interactions would rigidify the loop side chains and by extension produce clearer density. The modelled alanines likely do not accurately portray the potential for extensive interactions these residues may engage in with neighbouring N protomers. Indeed, residues homologous to these have been identified as putative drivers of clam-shell interactions in the paramyxoviruses NDV, NiV, and SeV [32,37,38]. Focussed refinement on this area failed to improve map resolution, and likely reflects – in part – the inherent flexibility of both this loop and the intercalating interface. Similarly, local resolution around the HeV N protein Ct-arm α17 was poor and while an alpha helix could confidently be built into the density, a number of side chains were also stubbed and modelled as alanines. Finally, more Ct-arm residues are resolved in the helical NiV N protein structure than in the HeV N double-ring structure, while the HeV N protein structure shows the very N-terminal Met1 residue where the NiV N protein structure cuts off at Ile4 [32]. Comparison of the cognate region in protein structures from related viruses revealed that the extreme N-terminal residues are either distended away towards the central cavity as in NDV [37] or are completely absent as for PIV5, NiV, MeV, and SeV [29,32,35,38] (Figure 8c-f). The shift in orientation or complete absence of the extreme N-terminal residues may be due to the presence of remaining N-terminal affinity tags that limit local cryoEM resolution. Indeed, an N-terminal affinity purification tag remained attached to resolved N protein structures for PIV5, NDV, NiV, and SeV [32,35,37,38] while the MeV purification used a C-terminal tag [29]. The tag-free N-terminal residues in the HeV double-ring assembly were identified as a potential minor interface (Figure 4). This elbow interface likely exists in NiV N protein assemblies as well, and the data suggest a similar interface may occur in more distantly related viruses such as PIV5 and MeV [29,35].

**Figure 8.**
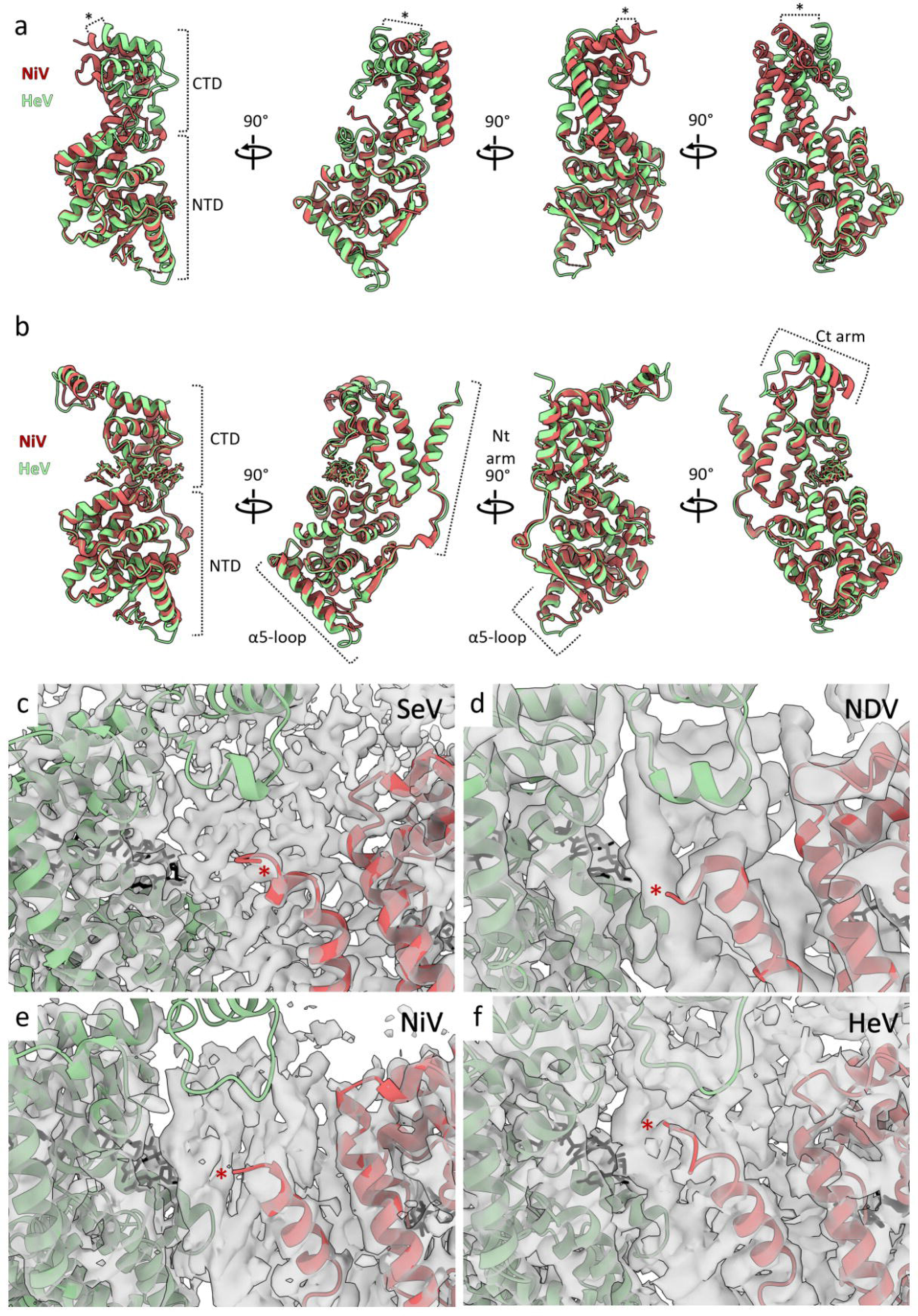
Comparison of the HeV N protein double-ring assembly to N protein assemblies from related viruses. **a-b**) Structural alignment of the HeV N protein model and two NiV N protein models. Models are presented as ribbons, with the HeV structure in green and the NiV structures in red. **a**) Alignment of the HeV N protein with the crystal structure of NiV N protein (PDB: 4CO6) with a global, pruned RMSD of 0.783 Å. The * indicates a 15.5 Å shift in position of the His369 Cα. The HeV N protomer-bound RNA hexamer was removed and the Nt- and Ct-arms were truncated to resemble the truncated apo-NiV N protein. The NiV P protein domain present in the crystal structure was also removed for clarity. **b**) Alignment of the HeV N protein with the helical cryoEM structure of NiV N protein (PDB: 7NT5) with a global, pruned RMSD of 0.756 Å. **c-f**) The structures of the multimeric N-protein assemblies of (**c**) SeV (PDB: 6M7D, EMD-30129), (**d**) NDV (PDB: 6JC3, EMD-9793, (**e**) NiV (PDB: 7NT5, EMD-12581), and (**f**) HeV (this study) are presented with the N_i+1_ (red) and N_i-1_ (green) protomers as ribbons and the bound RNA as black sticks inside their respective EM density maps (transparent grey). The N_i_ protomer models are not visualised for clarity but the density map is retained. The N_i+1_ utmost N-terminal residue resolved in each model is indicated by an asterisk, corresponding to SeV Gly3 (**c**), NDV Met1 (**d**), NiV Ile4 (**e**), and HeV Met1 (**f**) which are distended away from the N_i-1_ Ct-arm loop for SeV, NDV, and NiV (**c-e**) but not for HeV (**f**). The HeV N-terminal purification tag (hexahistidine-SUMO) was proteolytically removed prior to cryoEM grid preparation. The N-terminal hexahistidine-tag (SeV and NDV) or N-terminal hexahistidine-3C-tag (NiV) was retained but not resolved.

## Discussion

The emergence of novel pathogens such as HeV is accompanied by gaps in fundamental knowledge and technical resources. It is therefore of paramount importance to increase these key assets prior to an outbreak. While preparedness systems are falling into place for HeV, including surveillance, diagnostic tools and laboratory confirmation, and treatment and prevention, many questions regarding the molecular biology of HeV remain unanswered.

Following cryoEM data collection, a multimeric HeV N protein double-ring assembly was resolved to 3.5 Å resolution, enabling the building and refinement of an atomic HeV N protein model. Paramyxoviral double-ring complexes may represent a convergent architecture that reflects the need for 5’ capping of genomic and antigenomic nucleocapsids, the need to encapsidate leader sequences, and the need for short-term metastable N protein storage. The elucidation of the HeV N protein structure allowed for the identification of differences between it and related viral N protein structures, but importantly may also highlight commonalities that can be exploited for the design of new, broad-acting therapeutics.

The paramyxoviral nucleocapsid assembly entirely covers the viral genome, protecting it from detection and degradation by host factors and serving as the template for viral mRNA transcription and genome replication. Here, we determined the cryoEM structure of the recombinantly expressed, full-length and tag-free HeV N protein in complex with *E. coli* host-derived RNA. This structure is, to our best knowledge, the first high-resolution structure of the HeV N protein at 3.5 Å resolution and the first formal description of a paramyxoviral N protein double-ring assembly.

Overall, the HeV N protein fold resembles closely and more distantly related paramyxoviral N protein structures, including the recently solved N protein structures of PIV5 [35], MeV [29], NDV [37], and SeV [38], with its closest structural and genetic match being NiV N [32,44]. The N protein consists of an Ncore region and an intrinsically disordered Ntail, and, as is common for paramyxoviral N proteins, only the Ncore region was well resolved with no clear cryoEM density representing the Ntail. The HeV N protein shares around 92% amino acid identity with the NiV N protein [8,14,32,44]. The crystal structure for NiV N was solved as a truncated protein – missing both the Nt- and Ct-arms – and in complex with the Nt region of the NiV P protein [44]. More recently, the full NiV Ncore structure was solved through cryoEM from helical particles [32].

Structural alignment of the final HeV N protein model (PDB: 8CBW) with both the NiV N protein crystal structure (PDB: 4CO6) and cryoEM structure (PDB: 7NT5) revealed a globally similar protein fold, with some key differences. The most prominent of these differences is the approximately 25° rotation and up to 15.5 Å shift in the CTD, which is the result from the open, apo-configuration of the P protein-bound NiV N (P-N^0^) complex in the NiV N crystal structure [44]. Since the cryoEM NiV N structure is RNA-bound [32], it offers a better comparison with the similarly RNA-bound HeV N structure. A slight shift in the orientations of both the Nt- and Ct-arms likely stems from the differences in overall multimeric morphology, since the NiV structure originates from a helical segment with ∼13.2 N protomers per helical turn, while the HeV structure is drawn from a 14 N protomer-containing flat ring within the double-ring particle. Helical spiralling of a single, split tetradecameric ring would generate a helix resembling that of NiV and inevitably incur some minor conformational changes likely engendered by the flexible Nt- and Ct-arms whilst maintain a relatively static globular domain fold.

Furthermore, the NiV N Ct-arm has resolved four additional residues compared to the HeV N protein structure, while the HeV N protein structure resolves the extreme N-terminal Met residue where the NiV N protein structure terminates at Ile4 [32]. Similarly, the extreme N-terminal residues are often absent in N protein structures from other, related viruses, including PIV5, MeV, and SeV [29,35,38]. The resolution of the extreme N-terminal residues in the HeV N structure enabled the identification of a novel, albeit likely weak, interaction surface between the N_i+1_ and N_i-1_ protomers here called the elbow interface. The presence of an elbow interface in the N protein assemblies of PIV5, MeV, NiV, and SeV is likely obscured due to the absence of the extreme N-terminal residues in these structures. Where the expression and purification pipeline for HeV N included an affinity-tag removal step to generate a native protein sequence, those for the viral N proteins listed above did not. As a result, the presence of remaining, unstructured, N-terminal affinity tags may locally limit cryoEM resolution and fail to depict the elbow interface. This hypothesis is further supported by the N protein structure of NDV which does resolve the extreme N-terminal residues, but these face away from the putative elbow interface and towards the central cavity [37], likely due to the presence of an N-terminal affinity tag.

Finally, the solvent facing HeV NTD alpha-helix 5 (α5) and its succeeding loop show significant structural and sequence variability among paramyxoviruses and its position and length are highly variable among the N proteins of closely related paramyxoviruses, including MeV, NDV, NiV, PIV5, and SeV [29,32,35,37,38,44]. Compared to the cryoEM NiV N structure, the HeV α5-loop region contains two amino acid substitutions compared with the NiV N protein; L108V and E137D, both of which are solvent exposed [32]. In the context of a helical nucleocapsid, this solvent-exposed helix-loop region is likely important for interactions with both the RdRp complex and the M protein at various stages of the viral life cycle [17,48]. Furthermore, the α5-loop region is also part of the intercalating interface in the HeV double-ring assembly and in the clam-shell assemblies found for NDV, NiV, and SeV [32,37,38].

While the experimental particle stack only contained 5683 particles, D14 symmetry imposition resulted in an effective 157,864 asymmetric units. Symmetrised reconstructions often struggle to resolve regions of structural or compositional heterogeneity, and their signal may be diluted compared to the less heterogenic regions. In the reconstruction of the HeV N protein assembly, some of these heterogenic regions, foremost Met114-Asp119, are poorly resolved and as a result hindered confident model building of side chain residues. The electron density in the RNA binding cleft, identified through comparison with homologous structures, enabled *de novo* modelling of a hexameric RNA chain in the asymmetric unit. The electron density unambiguously identified the RNA backbone and the general location of the bases, but not their identities. This is to be expected due to both the presence of random host-derived RNA and symmetrised averaging of that RNA. Individual HeV N protein double-ring assemblies are unlikely to have complete occupancy by bacterial host-derived RNA over each of the 28 N protomers. Any RNA-free N protomers in the assembly would be outnumbered by RNA-bound protomers and as a result are averaged out against them, evidenced by the clear RNA density. Non-symmetrised particles would likely shed more light on both the uniformity of RNA occupancy as well as regions of structural flexibility with the requirement for larger particle stacks.

The RNA occupying the RNA binding cleft was modelled as one hexa-uridine per N protomer, adhering to the convention for paramyxoviral N proteins. The typical 3’ to 5’ ‘three in, three out’ base stacking orientation observed in other paramyxoviral N proteins is also found in the HeV N assembly, where bases 1, 2, and 3 face in towards the protein and bases 4, 5, and 6 are solvent exposed. Furthermore, a series of basic (Lys178, Arg192, Arg193, Arg352) and polar (Thr185, Gln199, Gln200, Tyr258, Gln319, Ser344, Tyr354) residues were identified as putatively interacting with the RNA chain through electrostatic and hydrophobic interactions. While most of these interactions are with the RNA backbone, a key, conserved interaction occurs between RNA bases 5 and 6 and the amino acid dyad Gln199 and Gln200 which HeV N shares with the N proteins from MeV, NiV, and PIV5 [29,32,35]. Additionally, a conserved in-base cavity is contiguous with the RNA binding cleft, consisting of two neighbouring N protomers and can accommodate larger purines in the in-base stack comprising nucleotides 1, 2, and 3. Putative interactions with residues Tyr258, Glu260, Glu261, Phe268, Lys 321, Ser344, and Met345 may facilitate purine binding where pyrimidines are too distant. A homologous cavity exists in closely related N protein assemblies, such as NiV, NDV, and MeV [29,32,37] and merits further investigation as a potential therapeutic target.

While N-RNA interactions are thought to be sequence non-specific and HeV N model building was done in a sequence-agnostic manner, some evidence for MeV suggests RNA encapsidation exhibits a preference for purine-rich sequences and importantly, the antigenomic leader sequence [49]. Similar preferences for purine rich and leader-specific RNA were also observed for the more distantly related RSV and VSV, respectively [50,51]. Indeed, the HeV genomic and antigenomic leader sequences are purine rich as well, with ∼70% and ∼63% purines for the genomic and antigenomic leaders, respectively. Moreover, the terminal six nucleotides of the HeV genomic and antigenomic leaders are identical (5’-ACCGAA) [14]. Additional interactions with these purine bases both in the in-base cavity as well as in the region of bases 5 and 6 may drive this difference in specificity and be important factors in the initial stages of 5’ encapsidation of the nascent genome as it emerges from the RdRp complex. As a result, the in-base cavity and the RNA binding cleft may present a target for the development of therapeutics that abrogate the critical step of RNA-encapsidation in the paramyxoviral life cycle.

HeV, discovered in 1994, is the type species for the genus *henipavirus* in the family *Paramyxoviridae* [52]. Emergent zoonotic outbreaks and surveillance of wild populations have expanded the genus with NiV, *Cedar henipavirus* (CedV), *Mojiang heninpavirus* (MojV), and *Ghanaian bat henipavirus* (GhV) [53–55] and over the past few years with the discovery of four new species; *Gamak henipavirus* (GAKV), *Daeryong henipavirus* (DARV), *Langya henipavirus* (LaV), and *Angavokely henipavirus* (AngV) [56–58]. The recent discovery of a novel HeV strain in Australia, called HeV-var [59], as well as the addition of four new henipaviral species in the past 3 years, highlights the needs for continued surveillance and research into this genus and HeV specifically. With around 88% sequence identity for the HeV-var N gene sequence compared to the HeV reference strain (NCBI RefSeq: NC_001906), a total of 18 amino acid substitutions were found of which 13 lie in the intrinsically disordered Ntail region (Supplementary Figure S8). Five substitutions are located in the Ncore region; S2G, D6E, and S13N in the Nt-arm, L153M in the NTD, and S386P in the Ct-arm (Supplementary Figure S8). Both D6E and S13N face the solvent of the double-ring central hole, however Ser2 forms part of the putative elbow interface between the N_i-1_ and N_i+1_ neighbours (Figure 4). The S2G substitution in HeV-var could abrogate this elbow interface. The NTD-located L153M substitution sits at the upper surface of the NTD facing the exterior solvent and is unlikely to be involved in intercalation zone interactions. Finally, the S386P substitution in the Ct-arm faces the solvent in the double-ring assembly, but likely forms part of the interface between successive helical turns in the nucleocapsid assembly. These interactions are thought to contribute minimally to nucleocapsid stability due to the nucleocapsid’s inherent flexibility and the observation that each helical turn consists of a non-integral number of N protomers [27]. Interestingly, no substitutions were described in the α5-loop region, which normally exhibits significant structural and sequence variability among paramyxoviruses. Taken together, the amino acid substitutions in the Ncore observed in the HeV variant strain likely have a minimal impact on HeV nucleocapsid formation and architecture. The large number of substitutions in the intrinsically disordered Ntail region, especially towards the C-terminus, could indicate a changed interaction with the RdRp complex and M proteins however these assumptions still await experimental validation.

The dimer of tetrameric rings that form the HeV N protein double-ring assembly is, to our best knowledge, a unique paramyxoviral N protein architecture that lacks a formal description in the current literature. However, other paramyxoviral N protein architectures have been observed that are reminiscent of the double-ring assembly. N protein clamshell interfaces have been described, in detail, for the paramyxoviruses NDV, NiV, and SeV NLFs [32,37,38]. In these cases, recombinant expression of N proteins revealed clamshell interfaces as both discrete particles reminiscent of, but distinct from, the double-ring assemblies [32,37] as well as part of a so-called double-headed helix; two anti-parallel NLFs stacked bottom-to-bottom and following the NLF’s helical register [37,38]. While a similar loop region is involved in these interactions in all three, the specifics of the clamshell architectures differ between the three reports. In the case of NDV, the N protomers that constitute the clamshell are positioned vertically in line and do not intercalate [37]. Those described for SeV appear to have two forms, called ‘closed’ and ‘hyper-closed’, where the level of intercalation is higher for the hyper-closed state [38]. Finally, the NiV clamshell exhibits extensive intercalation similar to the HeV double-ring assembly [32]. Furthermore, while in each of these cases the two faces of the clamshell follow the helical register of the NLF, an additional small population of NiV clamshells have been observed where the ∼13.2 N protomer helix is capped with a flat tetradecameric ring [32]. This ring closely matches one half of the HeV double-ring assembly. Importantly, the presence of double-headed helices was confirmed from virus-extracted nucleocapsids of SeV [38] and the inability to form double-headed nucleocapsids was detrimental to EGFP reporter gene expression in an NDV minigenome-based replicon assay [37].

The double decameric N protein rings of RSV, a member of the family *Pneumoviridae*, has recently been resolved through cryoEM [60]. In this RSV N protein assembly, akin to HeV, two N protein rings stack in a bottom-to-bottom orientation. Interestingly, the two rings do not intercalate to an appreciable degree and instead interact solely through the tips of the NTDs. The RSV N_i_ thus only interacts with one N protomer in the opposite ring (n_i_) where in the HeV N double-ring assembly the N_i_ protomer interacts with two N protomers (n_i_ and n_i+1_). As a consequence, the RSV double-ring assembly exhibits ten apertures along the interface between the two rings that made up of the NTDs of N_i_, N_i-1_, n_i_, and n_i+1_ which are over 13 Å wide at their narrowest point Interestingly, a recent paper describes the presence of single-faced closed-ring N protein assemblies inside RSV virions [36]. Although of lower resolution due to the use of cryo-ET, these rings bear a striking resemblance to the single and double decameric rings observed for recombinantly expressed RSV N protein [25,26,60] and indeed one tetradecameric half of the HeV double-ring assembly. Their abundance in RSV virions hints at a potential physiological relevance. While the occupancy and identity of RNA in these virion-enclosed rings was not confirmed, it is known that in RSV-infected cells short, 21 to 25 nucleotide-long RNAs generated from the genomic leader and trailer sequences are among the most abundant viral RNAs present [61–63]. Furthermore, some of these RNAs are resistant to nuclease treatment, indicating they may be encapsidated [62].

The occurrence of N protein double-rings and the related clamshells in recombinantly expressed N protein samples could be an artefactual result of the method of sample generation. However, the presence of clamshells in virion-extracted nucleocapsids [38] and of N protein ring assemblies inside virions [36] suggests otherwise. A picture emerges where the double-ring assembly may be indicative of one or multiple of a number of viral processes.

Firstly, it is hypothesised that the occurrence of clamshell interfaces both protects the viral genome from nuclease and protease attack and drives the inclusion of multiple copies of the viral genome into nascent virions [17,37,38]. Interestingly, these clamshells appear to join only the 5’ ends, which are the first to emerge from the RdRp during genome replication and may be the most vulnerable to degradation. Furthermore, polyploidy is increasingly recognised in a wide range of members of the order *Mononegavirales* [64–67] and clamshell interfaces may explain this occurrence at least partially. The double-ring resembles these clamshells, in particular the flat ring of the semi-spiral clamshell interface found in NiV N assemblies, and the double-ring assembly may therefore be a by-product of the viral need for genomic 5’ nucleocapsid capping.

Secondly, the high abundance of complementary leader and trailer RNAs in virus-infected cells could result in the activation of the innate immune response. The encapsidation of these sequences is hinted at for RSV [62], which could shield them from detection by the innate immune system. Interestingly, the 56 nt-long HeV leader sequence, which is transcribed and separated from the mRNA transcripts during the RdRp transcription mode, falls within the occupancy range for a single tetradecameric ring (84 nt). This purine-rich sequence is also likely preferentially encapsidated over RNA chains with an even purine-pyrimidine ratio [49–51]. The HeV trailer sequence is shorter than the leader and similarly rich in purine bases. An antagonistic function in stress granule formation and cellular apoptosis and has been reported for the (released) trailer sequence in SeV and RSV [68,69] and a similar function could exist for the HeV trailer. Whether putative encapsidation of the trailer aids or hinders this function is unclear, but encapsidation of both the leader and the trailer could permit inclusion into progeny virions and subsequent release in newly infected cells to perform their suggested antagonistic functions without immediate the requirement for viral RNA transcription.

Thirdly, single or double N-RNA rings such as the HeV double-ring assembly, could play a role in nucleating liquid-liquid phase separation for biomolecular condensates and aid in the formation of viral transcription/replication centres [70,71]. Evidence has been found for this process in RSV and has recently been investigated in detail for MuV where a combination of cryo-ET, fluorescence microscopy, and mass spectrometry reveal a stress-induced modulation of phase-separated N-P complexes [72]. A similar role may exist for the HeV double-ring assembly during phase separation in HeV infected cells.

Finally, the HeV N protein double-ring, and indeed N protein rings observed elsewhere, may reflect the need for short-term, reversible storage of N protomers in the absence of abundant P protein chaperones. Indeed, the P protein engages in extensive interactions with N protomers to maintain them in an open, apo-state (N^0^). Re-initiation of transcription by the RdRp complex is inefficient and as a result viral mRNA abundance is inversely correlated to the distance from the genomic 3’-end [17,73,74]. Consequently, the abundance of non-nucleocapsid-bound N protein is likely much higher than that of P protein and there is the potential for excessive N protein concentrations inside the infected cell. The formation of double-ring assemblies, either RNA-bound or in the apo-state, could therefore provide a short-term, metastable, multimeric N protein storage where P protein shuttles between them and the RdRp complex during active encapsidation of nascent genomic or antigenomic RNA.

## Materials and Methods

### Cloning *Hendra henipavirus* N gene into bacterial expression vectors

The HeV N gene sequence was taken from the reference *Hendra henipavirus* genome (NCBI Reference Sequence: NC_001906.3), codon-optimised for recombinant expression in *Escherichia coli* hosts, flanked by 5’ *Bam*HI and 3’ *Xho*I restriction sites, and supplied in pTwist (Twist Bioscience). Following digestion with *Bam*HI and *Xho*I, the restriction fragment was subcloned into pET-28a-6His-SUMO so that the HeV N protein coding region is downstream of the hexahistidine (6xHis) affinity tag and SUMO protease recognition sequence, with the resulting plasmid confirmed as correct by Sanger sequencing (Genewiz). The *Escherichia coli* (*E. coli*) DH5-α strain (Thermo Fisher Scientific) was used for plasmid amplification. Sequencing and amplification primers were designed using Benchling and ordered through Integrated DNA Technologies. Plasmid DNA was purified using a QIAprep Spin Miniprep kit (Qiagen) or QIAGEN Plasmid Midi kit (Qiagen) according to the manufacturer’s instructions. Where appropriate, products were resolved by agarose gel electrophoresis, excised from the gel and purified using a QIAquick Gel Extraction kit (Qiagen) according to the manufacturer’s instructions.

### Recombinant protein expression

*E. coli* BL21-Rosetta 2(DE3)pLysS (Novagen) transformed with pET-28a-6His-SUMO-HeVN was inoculated in a 1/500 ratio into 2xYT medium with kanamycin (16 gr/l Tryptone Plus (Sigma Aldrich), 10 gr/l Yeast Extract (Sigma Aldrich), 10 gr/l NaCl (Fisher Scientific), 50 μg/ml Kanamycin (Bio Basic)) and grown at 37°C in a Multitron Standard incubator (Infors HT) until reaching an OD_600_ of 0.5, at which point the growth temperature was reduced to 26°C and the inoculi were further grown until an OD_600_ of 0.8 was reached. Expression of HeV N was then induced by addition of 0.5 mM IPTG (ChemCruz) and cultures were further incubated for 4 hours at 26°C. Cultures were harvested by centrifugation for 25 minutes at 4500 rpm in an Avanti J26-XP with JLA 8.1000 rotor (Beckman Coulter).

Harvested pellets were resuspended in 30 ml lysis buffer (20 mM Tris-HCl pH 7.5 (ChemCruz), 500 mM NaCl, 20 mM Imidazole (Acros Organics), 0.1% v/v Triton X-100 (SAFC)) and subjected to one freeze-thaw cycle (−80°C to +40°C). The lysis buffer was then supplemented with cOmplete Mini Protease Inhibitor cocktail tablets (Sigma Aldrich), 10 μg/ml DNase I (Roche), 20 μg/ml RNase A (Invitrogen), and 0.25 μg/ml Lysozyme (Sigma Aldrich) and incubated for 30 minutes on ice with a magnetic stirrer. Lysates were sonicated for 10 cycles of 15 seconds on, 45 seconds off using the SoniPrep 150 Ultrasonic Disintegrator with External Process Timer (MSE), then transferred to ultracentrifuge tubes and centrifuged for 45 minutes at 16000 rpm and 4°C in an Evolution RC with SA300 rotor (Sorvall). The soluble fraction was collected in 50 ml falcon tubes and carried forward to purification.

### Affinity chromatography

Soluble fractions were passed through a primary HisTrap HP 5 ml column (GE Healthcare), previously equilibrated with column buffer (20 mM Tris-HCl pH 7.5, 500 mM NaCl, 20 mM Imidazole), using a 101U/R peristaltic pump (Watson Marlow). The column was washed with multiple washes of 25 ml column buffer with increasing concentrations of imidazole (20 mM, 50 mM, 100 mM, 300 mM, and 500 mM). The flow-through of the soluble fraction and each of the wash fractions was collected in full. The presence of protein was confirmed with Bradford Assay (Bio-Rad), NanoDrop One (Thermo Fischer Scientific), and SDS-PAGE analysis.

Fractions containing the majority of the Hev N protein were transferred into Spectra/Por 3 Standard RC Tubing (MWCO 3.5 kDa) (Spectrum). In-house recombinantly-expressed hexahistidine-tagged SUMO protease (Ulp1) was added to the dialysis tubing and fractions were dialysed overnight against dialysis buffer (20 mM Tris-HCl pH 7.5, 300 mM NaCl) at 4°C. SUMO protease cleavage was confirmed via SDS-PAGE analysis and the dialysed, cleaved mix was sterile filtered using a Minisart NML Syringe filter 0.45 μm SFCA (Sartorius) and passed through a secondary HisTrap column. The flow-through was collected and kept on ice, and the column was washed with pre-chilled low- and high-imidazole washes. The flow-through was sterile filtered and concentrated to a final volume of 7 ml using Amicon Ultra-15 Centrifugal filter concentrator (Merck Millipore).

### Size exclusion chromatography

Size exclusion chromatography (SEC) was performed using an ÄKTAprime Plus with HiPrep 26/60 Sephacryl 400 HR column (GE Healthcare) at 4°C, previously equilibrated with degassed and filtered SEC buffer (25 mM Tris-HCl pH 7.5, 100 mM NaCl). Fractions of 2.5 ml were collected following an 80 ml void volume. Presence of HeV N protein was confirmed with SDS-PAGE analysis. Concentrations and 260/280 ratios were determined and fractions containing HeV N protein were either directly used for electron microscopy (see below) or flash frozen and stored at −80°C.

### Preparation of cryo-grids

Copper Ultrathin Carbon (UC) Lacey grids (Agar Scientific) were glow discharged for 30 seconds at 10 mA on an easiGlow Glow Discharge Cleaning System. 3 μl protein sample at 1 mg/ml concentration was applied to the grids and plunge frozen into liquid ethane cooled in liquid nitrogen using an FEI Vitrobot Mark IV (Thermo Fisher Scientific; blot force 6, blot time 6-8s, humidity 95-100%).

### Data collection on cryo-grids

Data was collected on an FEI Titan Krios (X-FEG) TEM (Thermo Fisher Scientific) (ABSL, University of Leeds) operated at 300 keV with a Falcon III direct electron detector in integrating mode (Thermo Fisher Scientific), at a nominal magnification of 75000x and a pixel size of 1.065 Å. Further data acquisition parameters are shown in Supplementary Figure S2. 3548 micrographs were taken at a total dose of 59.7 e/Å^2^ split into 55 fractions and a defocus range of −0.7 to −2.8 μm. A detailed description of the data collection pipeline has been published [41]. Micrographs were visualised in ImageJ or RELION 3.1.1.

### Electron microscopy data processing and single particle analysis

The dataset of 3548 micrographs was collected using an automated approach and pre-processed on the fly during collection using the RELION software suite [41,42]. Motion correction and CTF estimation were carried out using RELION-integrated implementations of MotionCor2 (v1.2.1) [75] and Gctf (v1.18) [76], respectively. Following pre-processing, particles coordinates were manually selected using the manual picking job following the single-particle analysis (SPA) procedure. 3D maps were visualised in UCSF Chimera [77] and UCSF ChimeraX [78,79].

Using single particle analysis, a total of 3500 particles were selected from a random subset of 502 micrographs to generate a reference for automated picking through 2D classification. Reference-based automated picking was then optimised for picking threshold, maximum stddv noise, and minimum avg noise values, resulting in 875,262 selected particles, extracted using a box size of 360 pixels (383.4 Å). Multiple rounds of 2D classification were carried out to enrich the particle stack with true particles and removing junk particles. The high-resolution 2D classes revealed the presence of both a C2 and a C14 symmetry axis, for a combined D14 symmetry in the double-ring particle.

An initial 3D reference map was generated *de novo* by stochastic gradient descent (SGD) and used for 3D classification and 3D refinement. Manual re-alignment and resampling of the C1 symmetry refined 3D map to the grid of the unaligned map in UCSF Chimera [77] enabled D14 symmetry imposition in subsequent 3D jobs. The C1 symmetry refined 3D map was then used as a 3D reference map for subsequent D14 symmetry-imposed 3D classification and refinement jobs. A solvent mask was generated from the 3D class in Relion, using a lowpass filter of 15Å, an initial binarisation threshold of 0.012, a 3 pixel binary map extension, and a 3 pixel soft-edge. In post-processing, this solvent mask was applied to the refined cryoEM map to counter the influence of high-resolution noise to improve map quality and improve estimated resolution. The final EM map was deposited in the EMDB under accession EMD-16426.

### Atomic protein model refinement and interpretation

The cryoEM density map was of sufficient quality to allow the building and refinement of an atomic model. SWISS-MODEL [80] was used to predict an initial model based on homology search with PDB: 4CO6 [44], the closest match at the time of processing. The homology model was input for automated model building in Buccaneer [81]. The Buccaneer model was imported into Coot v0.9.4 [82,83] for manual refinement. High-resolution density enabled *de novo* building of a hexameric RNA chain modelled as poly-uridine following nucleoprotein structure convention. Multiple rounds of automated real-space refinement in Phenix v1.20.1 [84,85] and manual adjustment in Coot resulted in the final model. Manual adjustments were carried out on a single monomer in Coot which was then symmetry expanded in UCSF Chimera v1.15 according to D14 symmetry prior to real-space refinement to prevent erroneous refinement into non-cognate density. Poor local density abrogated confident side chain placement and forced the stubbing of some amino acids, which were modelled as alanines in the final model. Model validation was carried out as part of the final real-space refinement job through MolProbity [86,87] integrated within Phenix. Atomic models were deposited with the Protein Data Bank under accession 8C4H for the complete double-ring assembly and 8CBW for the asymmetric protomer unit. Interaction surfaces were calculated through PDBe PISA v1.52 [43] and 3D atomic models were visualised in UCSF ChimeraX v1.2.5 [78,79].

## Data Availability Statement

The refined cryoEM map was deposited with the EMDB under accession EMD-16426. Atomic models were deposited with the Protein Data Bank under accession 8C4H for the complete double-ring assembly and 8CBW for the asymmetric protomer unit.

## Acknowledgements

JNB, TAE and TCP were supported by funding from the European Union Horizon 2020 research and innovation programme, under the Marie Skłodowska-Curie Actions grant agreement no. 721367 (HONOURs). JBRW was supported by a Wellcome Trust 4-year PhD studentship (215064/Z/18/Z) and MJB was funded by grant BB/R00160X/1 from the BBSRC. The FEI Titan Krios microscopes were funded by the University of Leeds (UoL ABSL award) and The Wellcome Trust (108466/Z/15/Z). The FEI Vitrobot was funded by The Wellcome Trust, grant 218785/Z/19/Z. All *E. coli* strains were produced in house by Susan Matthews. Valuable suggestions during cryoEM data processing and model building were given by Dr. David Nicholson and Dr. David Klebl.

## Author Contributions

TCP, JNB, and TAE conceptualised the work. JNB and TAE were responsible for primary supervision of the investigation. TCP carried out the investigation. DPM assisted in cryoEM data collection. JBRW and MJB provided critical input in data processing. TCP visualised the data and prepared the manuscript. JNB, TAE, and NAR provided valuable feedback on the investigation and manuscript. JNB and TAE acquired funding for the work.

