## Supplementary material for "The cryoEM structure of the *Hendra henipavirus* nucleoprotein reveals insights into paramyxoviral nucleocapsid architectures": combined supplemental data (from another version of this preprint: version 1, 2023-12-07)

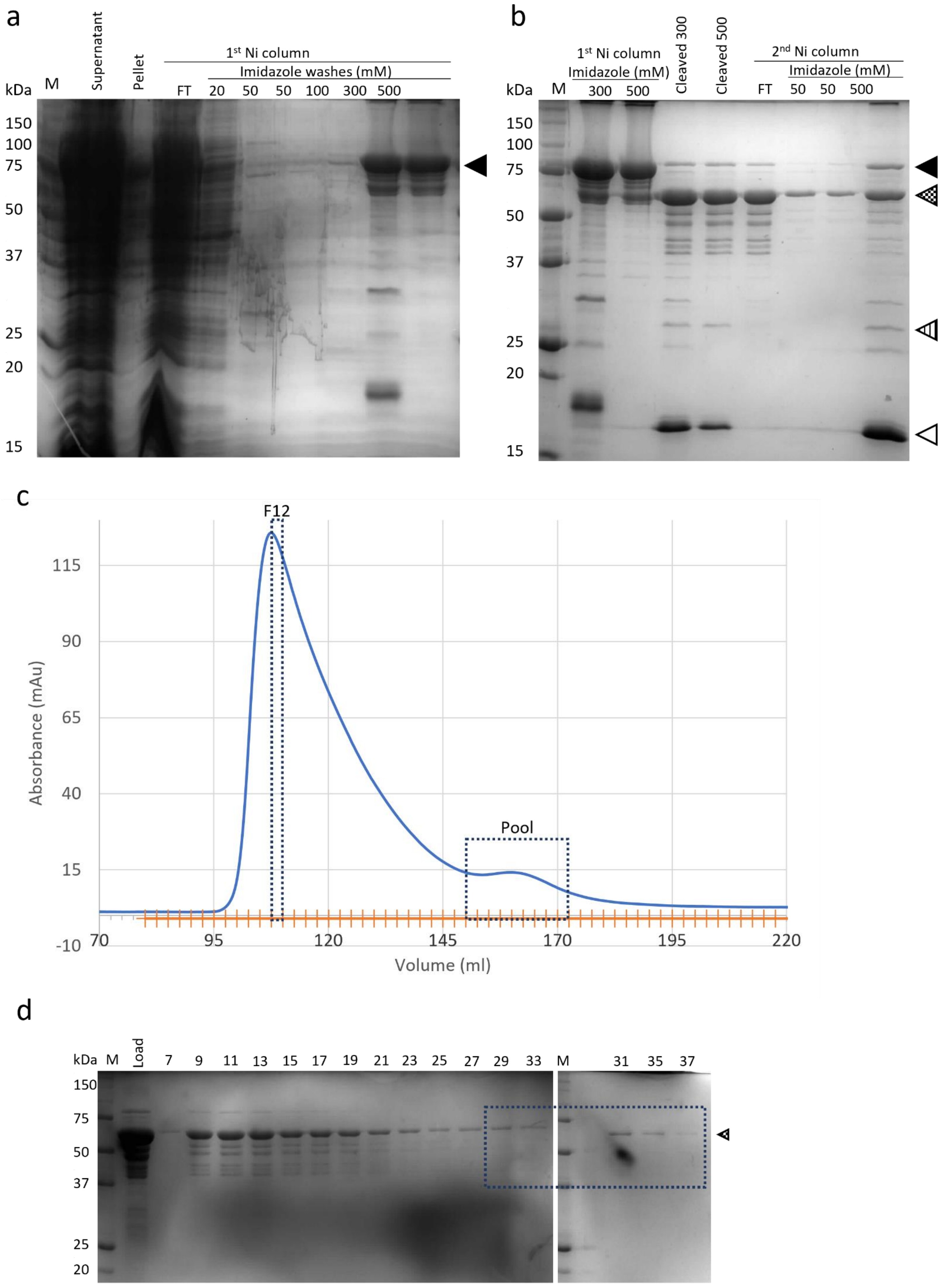
a
kDa
M
Supernatant
Pellet
1st Ni column
Imidazole washes (mM)
FT
20
50
50
100
300
500
150
100
75
50
37
25
20
15
b
kDa
M
1st Ni column
Imidazole (mM)
300
500
Cleaved 300
Cleaved 500
2nd Ni column
Imidazole (mM)
FT
50
50
500
150
100
75
50
37
25
20
15
c
F12
Pool
Absorbance (mAu)
115
90
65
40
15
-10
70
95
120
145
170
195
220
Volume (ml)
d
kDa
M
Load
7
9
11
13
15
17
19
21
23
25
27
29
33
M
31
35
37
150
75
50
37
25
20

**Supplementary Figure S1. Recombinant HeV N expression and purification.**

Coomassie stained SDS-PAGE of expression and purification samples. **a**) Fractions from expression and primary HisTrap column show a large portion of soluble HeV N protein (supernatant). The soluble HeV N was retained well in the HisTrap column and eluted, as expected, at 300 mM and 500 mM imidazole. **b**) Fractions from secondary HisTrap column, post SUMO protease treatment of HeV N. Bands indicate successful proteolytic cleavage of the 6His-SUMO HeV N fusion protein. Black arrowheads indicate 6His-SUMO HeV N fusion protein. Checkered arrowheads indicate native HeV N protein. White arrowheads indicate the post-cleavage 6His-SUMO tag. Striped arrowheads indicate 6His-tagged SUMO protease (27 kDa). **c**) Size exclusion chromatography (SEC) UV trace from a Sephacryl S400 26/600 column. Continuous UV absorbance readings are plotted in blue. Automatically collected 2.5 ml fractions are indicated by the orange hashed line. Fraction 12 (F12) and Pooled fractions (Pool) are outlined. **d**) Coomassie stained SDS-PAGE showing the presence of HeV N protein. The lane marked Load represents the sample loaded into the SEC column for reference. The numbered lanes represent SEC fractions loaded on gel and correspond to those in panel **c**. Chequered arrowheads indicate native HeV N protein.

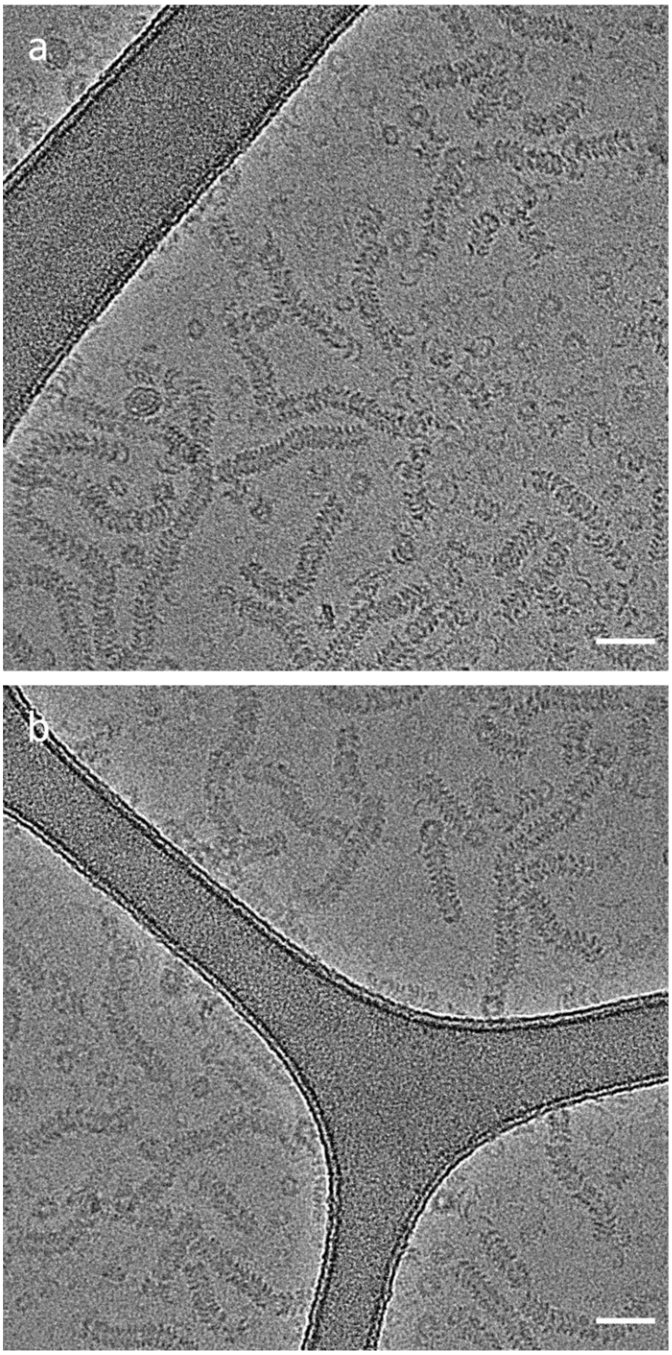


c

| Hardware | |
|---|---|
| Microscope | Krios 1 |
| Detector (mode) | Falcon III (Integrating) |
| Accelerating voltage (keV) | 300 |
| Pixel size (Å) | 1.065 |
| **Data acquisition parameters** | |
| Nominal magnification | 75k |
| Spot size | 4 |
| Illuminated area | 1.35 |
| **Dose** | |
| Square pixel (Å$^2$) | 1.13 |
| Dose per physical pixel per second | 45 |
| Dose per Å$^2$/sec | 39.8 |
| Exposure time (seconds) | 1.5 |
| Total dose (e/Å$^2$) | 59.7 |
| Number of fractions | 55 |
| Dose per fraction (e/Å$^2$) | 1.1 |
| **EPU parameters** | |
| Defocus range (-µm) | -0.7 to -2.8 (step -0.3) |
| Autofocus | After distance 10 µm |
| Drift measurement | N/A |
| Delay after stage shift | 5 sec |
| Delay after image shift | 0 sec |
| Exposures per hole | 1 |
| **Apertures (size in microns)** | |
| C1 | 2000 |
| C2 | 70 |
| C3 | 2000 |
| Objective | 100 |
| **General Information** | |
| Physical pixel size (Falcon III) | 14 µm |
| Cs | 2.7 |
| Micrographs taken | 3548 |

**Supplementary Figure S2. Data collection on UC Lacey grid.**
**a**-**b**) Representative micrographs from the FEI Titan Krios automated data collection through EPU, depicting HeV RLFs of all three predominant morphologies. Scale bars are 50 nm. **c**) Automated data collection parameters.

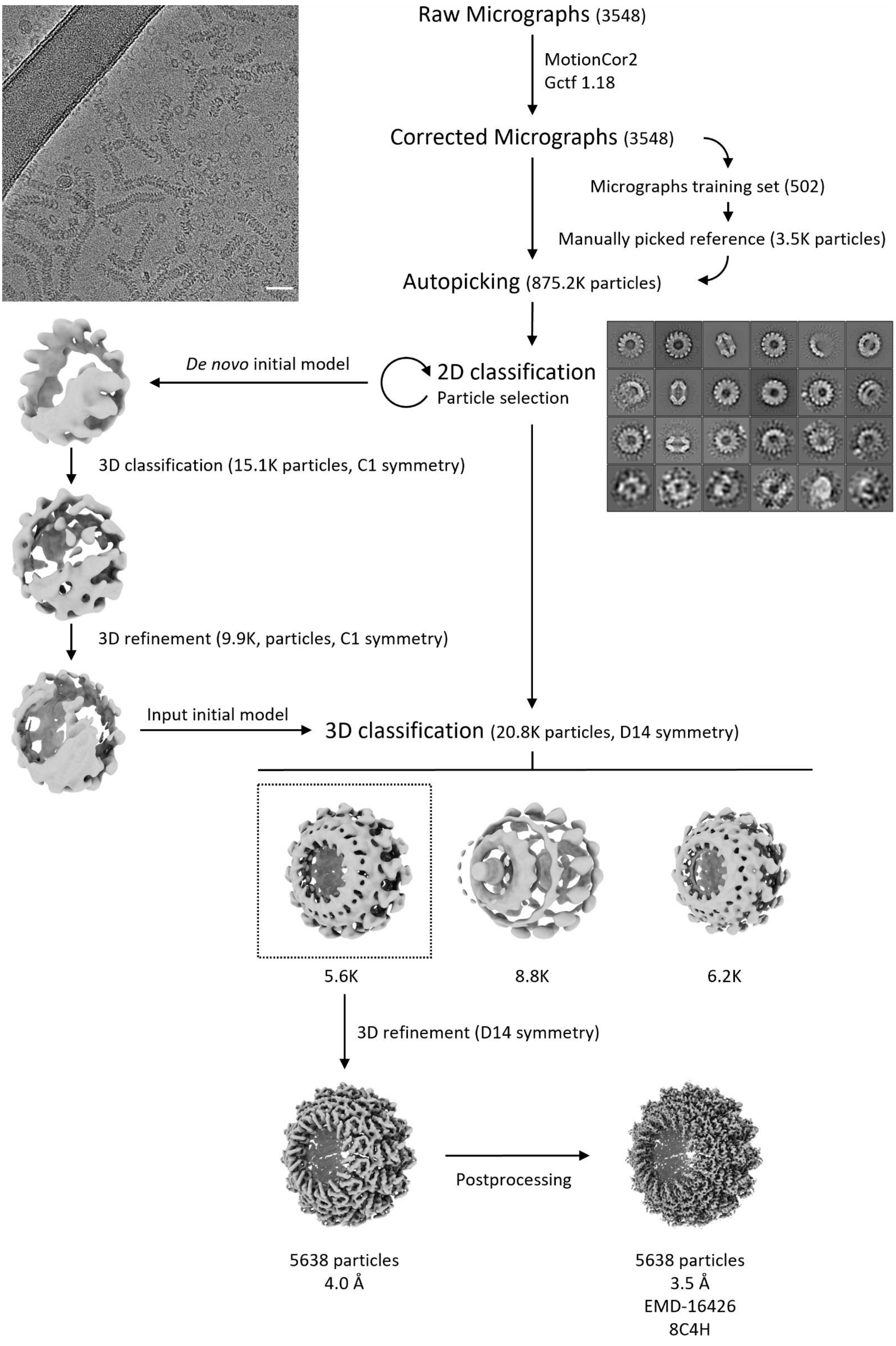
Raw Micrographs (3548)
MotionCor2
Gctf 1.18
Corrected Micrographs (3548)
Micrographs training set (502)
Manually picked reference (3.5K particles)
Autopicking (875.2K particles)
2D classification
Particle selection
De novo initial model
3D classification (15.1K particles, C1 symmetry)
3D refinement (9.9K, particles, C1 symmetry)
Input initial model
3D classification (20.8K particles, D14 symmetry)
5.6K
8.8K
6.2K
3D refinement (D14 symmetry)
Postprocessing
5638 particles
4.0 Å
5638 particles
3.5 Å
EMD-16426
8C4H

**Supplementary Figure S3. CryoEM data processing workflow.**

Following motion correction and CTF estimation of 3548 micrographs, 3500 double-ring particles were manually picked in Relion 3.1.1 from a subset of 502 micrographs and used as a reference for automated picking. Over multiple rounds of 2D classification and particle selection, a curated set of 20,809 particles was generated and taken forward for 3D classification. A 3D initial reference model was generated *de novo* and D14 symmetry was imposed on the 3D classes. 3D refinement of the optimal class (dashed box) resulted in a 4.0 Å EM map. Postprocessing using a mask generated from the 3D refined map then generated the final cryoEM map at 3.5 Å resolution, which was deposited to the EMDB under accession number EMD-16426 and the resulting structure was deposited to the PDB under accession number 8C4H.

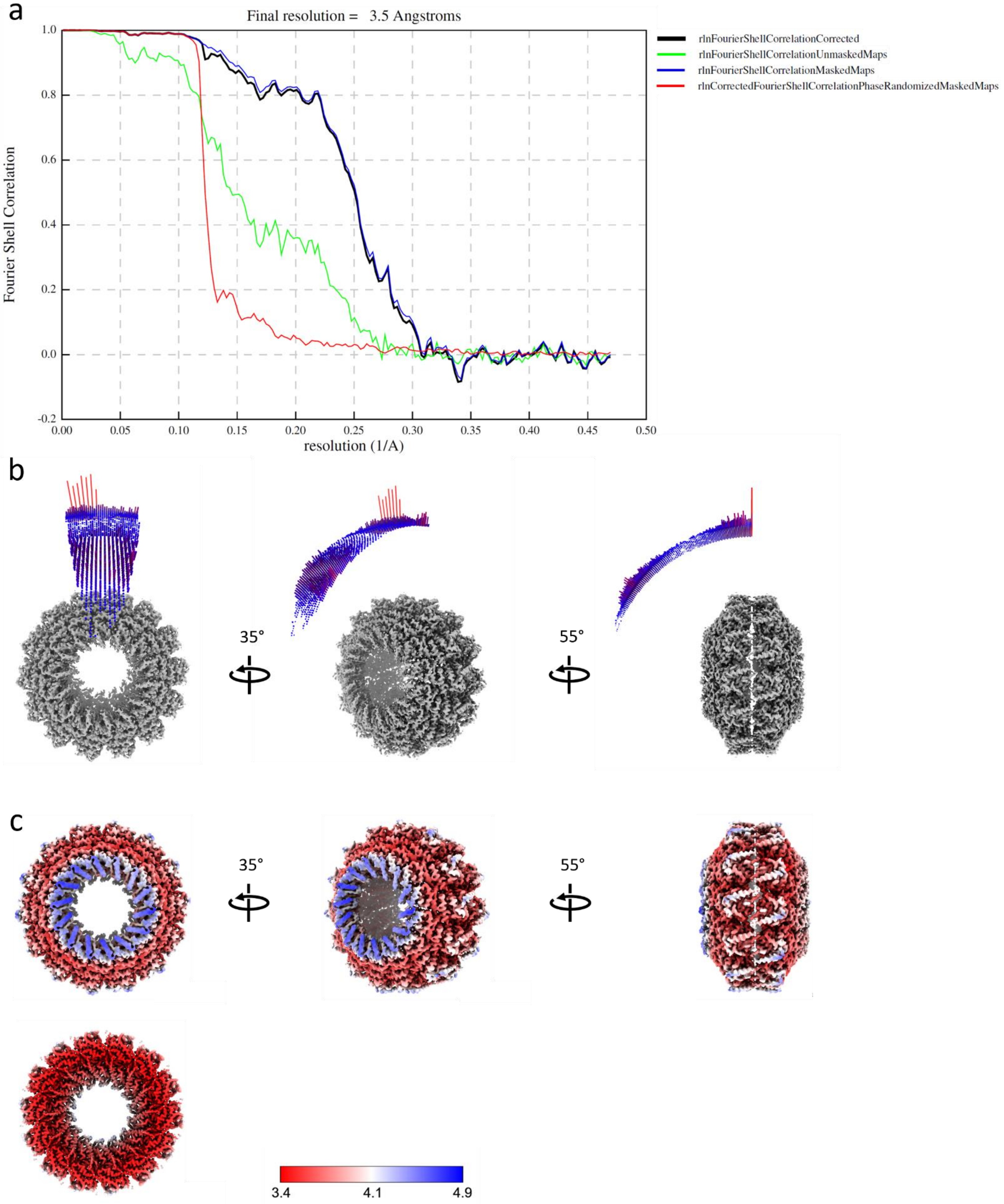


**Supplementary Figure S4. Symmetry refinement and post-processing for the HeV double-ring map.**
**a**) Fourier shell correlation plot from post-processing of the initial cryoEM map refinement. **b**) Angular distribution plot for the post-processed cryoEM map. Angular distributions appear in a wedge shape due to the imposed D14 symmetry. **c**) Local resolution representation of the HeV double-ring map. Colours range from 3.4 Å to 4.9 Å.

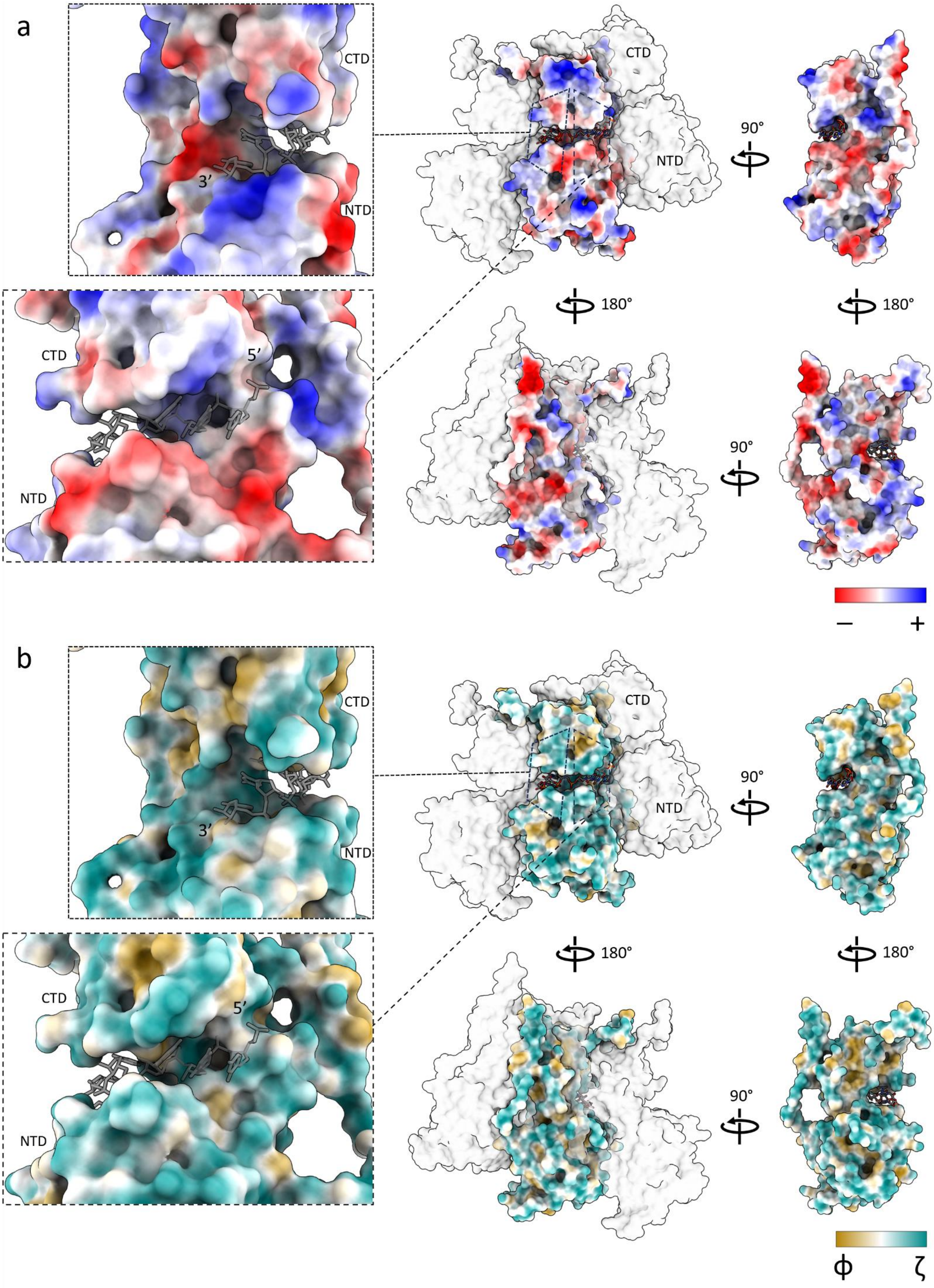

a
CTD
3′
NTD
CTD
NTD
90°
180°
180°
CTD
5′
NTD
90°
–
+
b
CTD
3′
NTD
CTD
NTD
90°
180°
180°
CTD
5′
NTD
90°
ϕ
ζ

**Supplementary Figure S5. HeV N protomer surface values.**

The HeV N protomer, with RNA (colour by atom in the overviews or transparent grey in the insets), has been coloured according to electrostatic potential (**a**) and hydrophobicity (**b**). The NTD and CTD as well as the RNA 3’ and 5’ ends (in the insets) are indicated. **a**) Electrostatic surface potentials are coloured for negative (red, -) and positive (blue, +) residues or uncoloured (white) for neutral residues. **b**) Colours are mapped according to Kyte-Doolittle hydropathy [88] from most hydrophilic (green, ζ) to most hydrophobic (gold, ϕ).

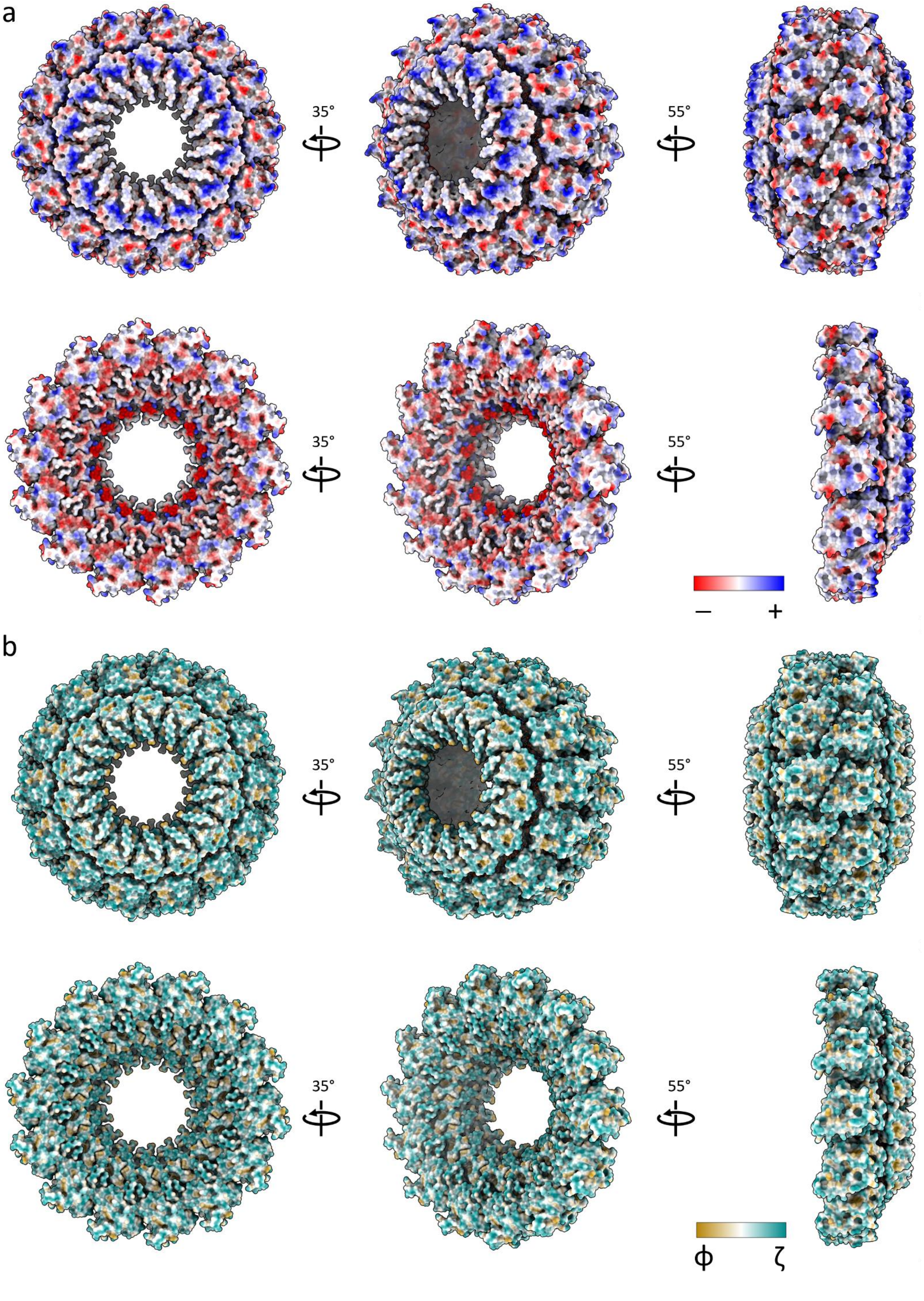

a
35°
55°
35°
55°
–
+
b
35°
55°
35°
55°
ϕ
ζ

**Supplementary Figure S6. HeV N assembly surface values.**

The HeV N protein double-ring assembly, with RNA, has been coloured according to electrostatic potential (**a**) and hydrophobicity (**b**). Top panels depict the dimer of tetradecameric rings. Bottom panels depict a single tetradecameric ring. **a**) Electrostatic surface potentials are coloured for negative (red, -) and positive (blue, +) residues or uncoloured (white) for neutral residues. **b**) Colours are mapped according to Kyte-Doolittle hydropathy [88] from most hydrophilic (green, ζ) to most hydrophobic (gold, ϕ).

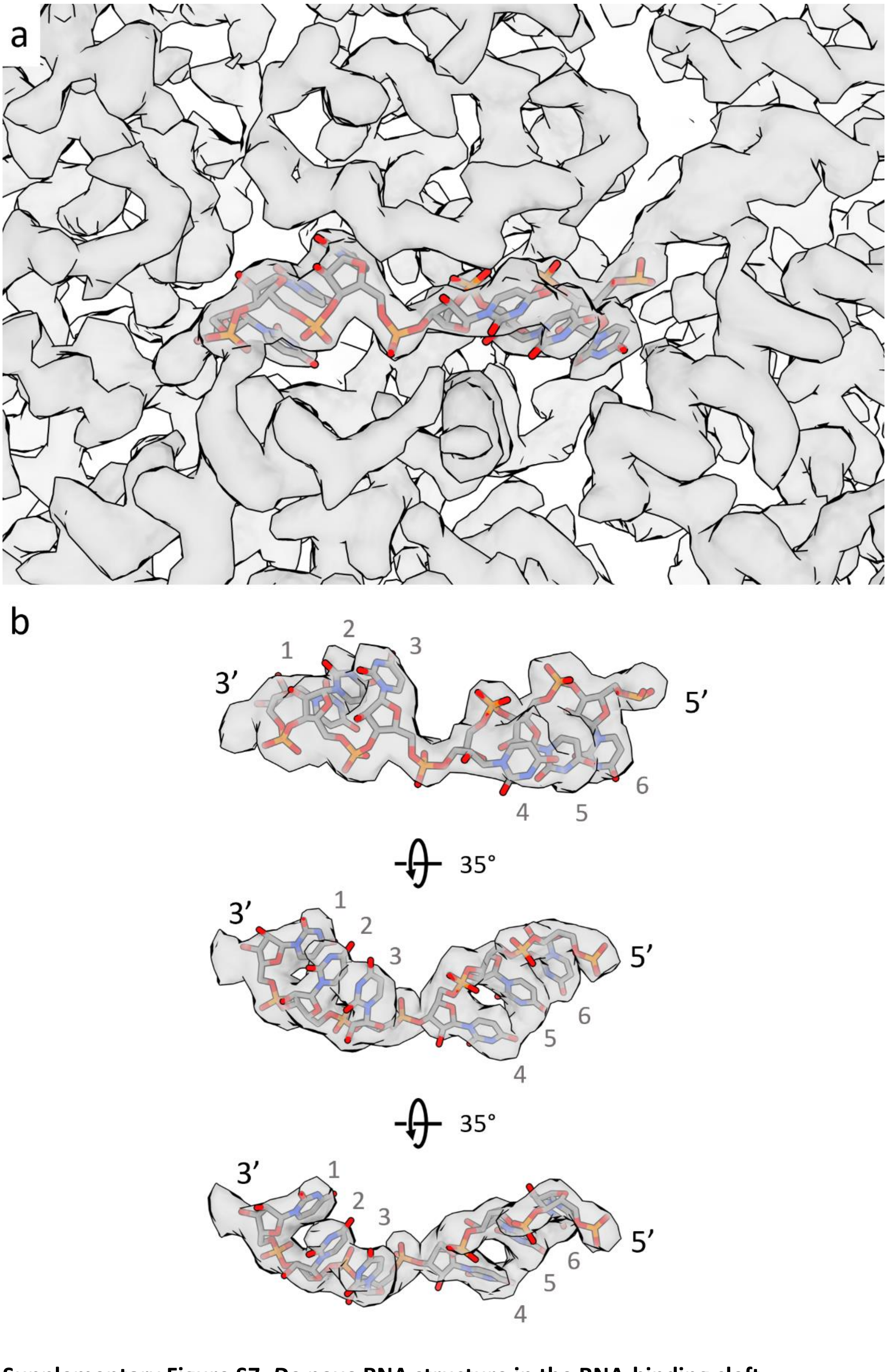


**Supplementary Figure S7.** ***De novo*** **RNA structure in the RNA-binding cleft.**
**a**) CryoEM density surrounding the N protein RNA-binding cleft depicting the *de novo* built hexa-uridine chain (sticks). **b**) The density occupying the RNA-binding cleft was isolated from the map and shows the built hexa-uridine RNA chain. The RNA 3’ and 5’ ends are indicated. Uridine nucleotides are numbered from the 3’ end and exhibit two characteristic base stacks, one facing in towards the N protein density (bases 1-3) and one facing out towards the solvent (bases 4-6).

a

```
HeV_ref MSDIFDEAAS FRSYQSKLGR DGRASAATAT LTTKIRIFVP ATNSPELRWE LTLFALDVIR SPSAAESMKI GAAFTLISMY SERPGALIRS LLNDPDIEAV IIDVGSMLNG 110
HeV_var MGDIFEEAAS FRNYQSKLGR DGRASAATAT LTTKIRIFVP ATNSPELRWE LTLFALDVIR SPSAAESMKI GAAFTLISMY SERPGALIRS LLNDPDIEAV IIDVGSMLNG 110
        *.***:**** **.******* ********** ********** ********** ********** ********** ********** ********** ********** **********

HeV_ref IPVMERRGDK AQEEMEGLMR ILKTARESSK GKTPFVDSRA YGLRITDMST LVSAVITIEA QIWILIAKAV TAPDTAEESE TRRWAKYVQQ KRVNPFFALT QQWLTEMRNL 220
HeV_var IPVMERRGDK AQEEMEGLMR ILKTARESSK GKTPFVDSRA YGMRITDMST LVSAVITIEA QIWILIAKAV TAPDTAEESE TRRWAKYVQQ KRVNPFFALT QQWLTEMRNL 220
        ********** ********** ********** ********** **:******* ********** ********** ********** ********** ********** **********

HeV_ref LSQSLSVRKF MVEILMEVKK GGSAKGRAVE IISDIGNYVE ETGMAGFFAT IRFGLETRYP ALALNEFQSD LNTIKGLMLL YREIGPRAPY MVLLEESIQT KFAPGGYPLL 330
HeV_var LSQSLSVRKF MVEILMEVKK GGSAKGRAVE IISDIGNYVE ETGMAGFFAT IRFGLETRYP ALALNEFQSD LNTIKGLMLL YREIGPRAPY MVLLEESIQT KFAPGGYPLL 330
        ********** ********** ********** ********** ********** ********** ********** ********** ********** ********** **********

HeV_ref WSFAMGVATT IDRSMGALNI NRGYLEPMYF RLGQKSARHH AGGIDQNMAN KLGLNSDQVA ELAAAVQETS VGRQDNNMQA REAKFAAGGV LVGGGEQDID EEEEPIEHSG 440
HeV_var WSFAMGVATT IDRSMGALNI NRGYLEPMYF RLGQKSARHH AGGIDQNMAN KLGLNPDQVA ELAAAVQETS VGRQDNNMQA REAKFAAGGV LVGGGEQDID EDEEPIEHSG 440
        ********** ********** ********** ********** ********** ***** **** ********** ********** ********** ********** *:********

HeV_ref RQSVTFKREM SMSSLADSVP SSSVSTSGGT RLTNSLLNLR SRLAAKAIKE STAQSSSERN PPNNRPQADS GRKDDQEPKP AQNDLDFVRA DV- 532
HeV_var RQSVTFKREM SMSSLADSLP SSSVSTSGGT RLTNSLLNLR SRLAAKAMKD NAAQSSVEKN TPNNRPQADS RGKEDQESKP AQNDLDFVRA DV- 532
        ********** ********:* ********** ********** *******:*: .:**** *:*  *********   *:*** ** ********** **
```

b

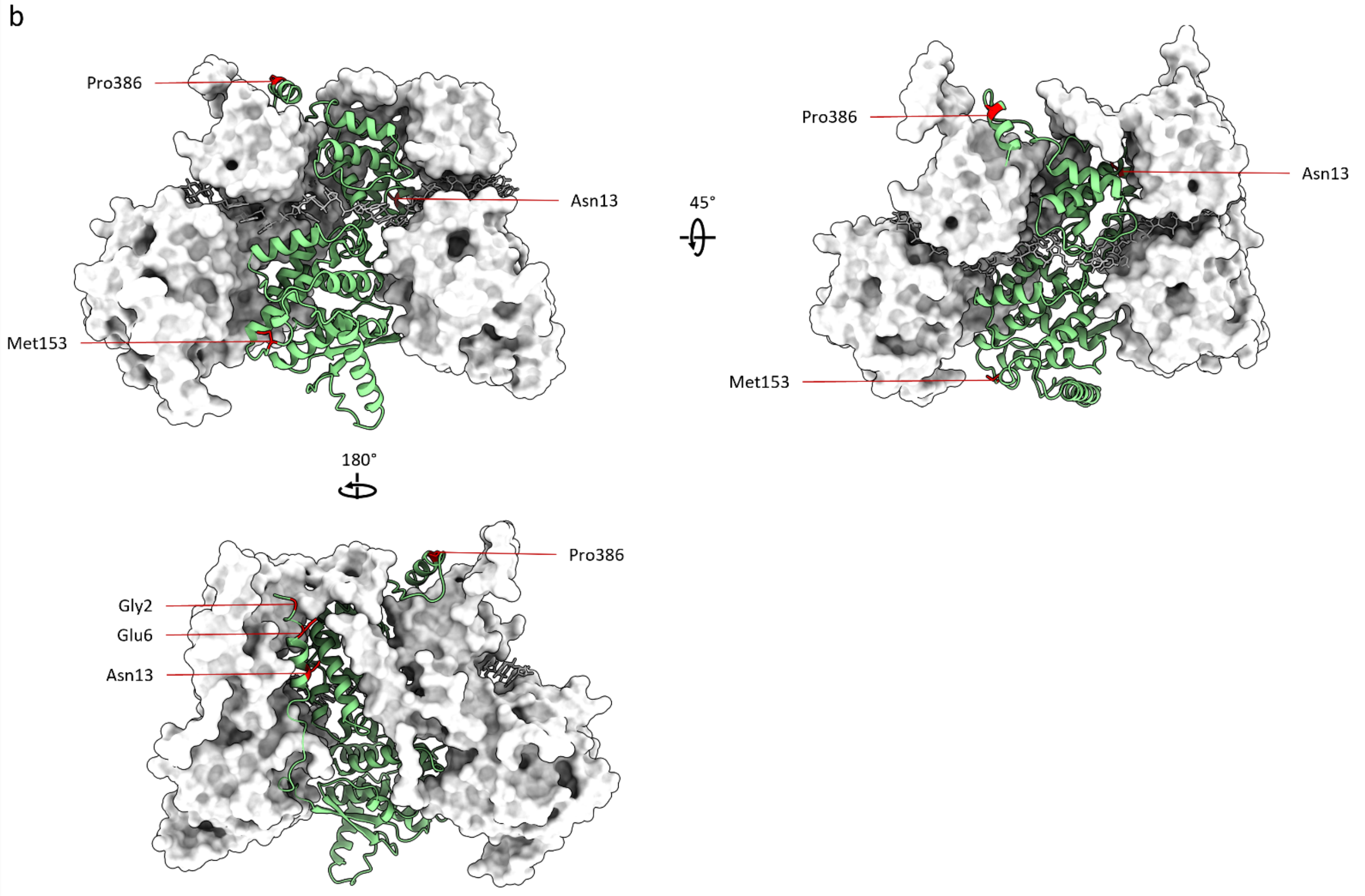


**Supplementary Figure S8. Mapping HeV variant amino acid substitutions.**

**a**) The full-length nucleoprotein sequences of HeV (HeV_reference; NCBI NC_001906.3) and HeV-var (HeV_variant; GenBank MZ318101) were aligned in Clustal Omega. Amino acid substitutions are indicated in red. N protein domains are indicated for Ncore; Nt-arm (blue), NTD (orange), CTD (green), Ct-arm (yellow), and the Ntail (grey). **b**) Amino acid substitutions between the HeV reference strain and the HeV variant strain are mapped (red) onto a single HeV N protomer (lime) flanked by two N protomers (white) in the assembled ring. The $N_i$ protomer (lime) is visualised as ribbons and the $N_{i-1}$ and $N_{i+1}$ protomers are surface rendered (white). Residues present in the HeV variant strain are indicated.

**Supplementary Table S1. HeV N protein double-ring assembly model validation**

| | |
|---|---|
| **Data processing** | |
| Symmetry imposed | D14 |
| True particle images (no.) | 5638 |
| Symmetrised particle images (no.) | 157,864 |
| Estimated map resolution (Å) | 3.48545 |
| FSC threshold | 0.143 |
| Map sharpening B factor ($Å^2$) | -132.352 |
| **Model composition** | |
| Non-hydrogen atoms | 87752 |
| Protein residues | 11032 |
| Nucleic acid residues | 168 |
| **Model validation** | |
| RMS Bond lengths (Å) | 0.002 (0) |
| RMS Bond angles (°) | 0.502 (0) |
| MolProbity score | 1.72 |
| Clash score | 9.57 |
| Rotamer outlier (%) | 0.13 |
| Ramachandran Favoured (%) | 96.63 |
| Ramachandran Outliers (%) | 0.00 |
| **Rama-Z (Ramachandran plot Z-score, RMSD)** | |
| whole (N=10976) | 0.80 (0.08) |
| helix (N=6524) | 1.95 (0.06) |
| sheet (N=504) | -0.11 (0.22) |
| loop (N=3948) | -1.95 (0.08) |
| Cβ outliers (%) | 0.00 |
| **Peptide plane (%)** | |
| Cis proline/general | 0.0/0.0 |
| Twisted proline/general | 0.0/0.0 |
| CaBLAM outliers (%) | 3.68 |
| **ADP (B-factors)** | |
| Iso/Aniso (#) | 87752/0 |
| Protein (min/max/mean) | 14.79/167.80/65.80 |
| Nucleotide (min/max/mean) | 52.67/104.15/71.05 |
| Ligand (min/max/mean) | --- |
| Water (min/max/mean) | --- |
| **Occupancy** | |
| Mean | 1.00 |
| occ = 1 (%) | 100.00 |
| 0 < occ < 1 (%) | 0.00 |
| occ > 1 (%) | 0.00 |
